# A physicochemical model of odor sampling

**DOI:** 10.1101/2020.06.16.154807

**Authors:** Mitchell E. Gronowitz, Adam Liu, Qiang Qiu, C. Ron Yu, Thomas A. Cleland

**Affiliations:** Department of Psychology, Cornell University Ithaca, NY 14853; Stowers Institute for Medical Research, 1000 East 50^th^ Street, Kansas City, MO 64110; Department of Anatomy and Cell Biology, University of Kansas Medical Center, 3901 Rainbow Boulevard, Kansas City, MO 64110

**Author notes:** Corresponding author: Thomas A. Cleland, Dept. Psychology, 278E Uris Hall, Cornell University, Ithaca, NY 14853, USA.

**Keywords:** Q-space, olfactory white, occupancy, pharmacology, binding, antagonist, Kir2.1, olfaction

## Abstract

We present a general physicochemical sampling model for olfaction, based on established pharmacological laws, in which arbitrary combinations of odorant ligands and receptors can be generated and their individual and collective effects on odor representations and olfactory performance measured. Individual odor ligands exhibit receptor-specific affinities and efficacies; that is, they may bind strongly or weakly to a given receptor, and can act as strong agonists, weak agonists, partial agonists, or antagonists. Ligands interacting with common receptors compete with one another for dwell time; these competitive interactions appropriately simulate the degeneracy that fundamentally defines the capacities and limitations of odorant sampling. The outcome of these competing ligand-receptor interactions yields a pattern of receptor activation levels, thereafter mapped to glomerular presynaptic activation levels based on the convergence of sensory neuron axons. The metric of greatest interest is the mean discrimination sensitivity, a measure of how effectively the olfactory system at this level is able to recognize a small change in the physicochemical quality of a stimulus.

This model presents several significant outcomes, both expected and surprising. First, adding additional receptors reliably improves the system’s discrimination sensitivity. Second, in contrast, adding additional ligands to an odor scene initially can improve discrimination sensitivity, but eventually will reduce it as the number of ligands increases. Third, the presence of antagonistic ligand-receptor interactions produced clear benefits for sensory system performance, generating higher absolute discrimination sensitivities and increasing the numbers of competing ligands that could be present before discrimination sensitivity began to be impaired. Finally, the model correctly reflects and explains the modest reduction in odor discrimination sensitivity exhibited by transgenic mice in which the specificity of glomerular targeting by primary olfactory neurons is partially disrupted.

**Author Summary:** We understand most sensory systems by comparing the responses of the system against objective external physical measurements. For example, we know that our ability to distinguish small changes in color is greater for some colors than for others, and that we can distinguish sounds more acutely when they are within the range of pitches used for speech. Similar principles presumably apply to the sense of smell, but odorous chemicals are harder to physically quantify than light or sound because they cannot be organized in terms of a straightforward physical variable like wavelength or frequency. That said, the physical properties of interactions between chemicals and cellular receptors (such as those in the olfactory system) are well understood. What we lack is a systematic framework within which these pharmacological principles can be organized to study odor sampling in the way that we have long studied visual and auditory sampling. We here propose and describe such a framework for odor sampling, and show that it successfully replicates some established but unexplained experimental results.

## Introduction

Unlike the visual and auditory modalities, the environmental sampling metrics of olfaction remain unclear. That is, whereas olfactory sensory transduction *per se* is straightforward, there is no underlying metric of physical similarity akin to chromatic wavelength, retinotopic spatial proximity, or auditory pitch to provide a quantitative external basis for odor similarity. Instead, odor similarity has been mapped heuristically [1, 2], with homologous series of simple molecules and variable-ratio binary mixtures serving as behaviorally validated sets of locally sequentially similar odorants. Indeed, in lieu of externally defined axes of similarity, these heuristic odor series have been invaluable for exploration of the similarity-dependent computations of the early olfactory system [1, 3–9]. It is reasonably clear that small differences in the activation profile across an animal’s complement of odorant receptors (~1000 types in mice and rats; ~400 in humans) underlie similar odor percepts whereas larger differences underlie more dissimilar percepts [1, 10, 11], particularly if one neglects the substantial effects of odor learning on perceived similarities [5, 12–14]. However, the transformation between the physicochemical space of an olfactory scene and the activation profile of the primary odorant receptors expressed by an animal sampling that scene has remained essentially undefined.

For many purposes in olfactory neuroscience, this transformation is unimportant. Experimental and theoretical studies of odor similarity spaces and their transformations in the neural circuitry of the early olfactory system require only the activation profiles of primary odorant receptors, or of the olfactory bulb glomeruli on which they converge; these receptor profiles define an organism-dependent olfactory metric space that we term R-space (see *Discussion*) into which similarity can be mapped. The details of sampling and transduction that underlie a given receptor activation profile are academic in this context, because by definition any physicochemical information that does not affect the activation of primary sensory neurons is not incorporated into sensory processing, and the sensory response has no means of detecting whether receptor activation is due to one ligand species alone, or a cocktail of multiple ligand species (such as comprise most natural odors). However, there are other scientific questions that depend on a theoretical understanding of the physicochemical sampling transformation. For example, understanding “active sampling” behaviors requires exploration of the underlying physicochemical sampling challenges as well as organismal goals [15–17]. Sampling studies of natural visual scenes have revealed that animals’ visual systems are adapted to the spatial frequency statistics of the natural world [18–20]. Many practical applications of research into sensory systems prescribe modifications of the physicochemical environment, necessitating some predictive understanding of these modifications’ effects on sensory systems. Accordingly, we sought to develop a general theory of physicochemical sampling, by constructing a single consistent framework, arising from first principles, to describe how olfactory systems sample the chemical features of environments.

To date, efforts to describe olfactory physicochemical sampling have been largely heuristic. The primary strategy has been to compose lists of many physicochemical properties, or molecular descriptors, of various odorous molecules and cross-reference these lists with either perceptual datasets [21–24] or the physiological response profiles of olfactory bulb glomerular arrays [25, 26]. While significant relationships with perceptual similarity certainly can be found, and their principal components can be labeled (e.g., pleasantness, directed dipole, ease of electron promotion; [21, 22]), this is not ultimately a generative process. To wit, it cannot be used to model the decomposition of odor mixtures [27], nor to strongly predict the activity of bulbar principal neurons [28], nor to study how the same physicochemical odorscene may be differently sampled by the noses of different animal species (potentially resulting in different patterns of similarity within their respective R-spaces). Moreover, the use of molecular descriptors is less concrete than it may initially seem, as the aggregate physical properties of the whole molecule gloss over the intramolecular heterogeneities such as localized charge distributions that directly influence ligand-receptor interactions (notably, single odorant molecules present multiple prospective binding sites to different receptors, the conformation of which is critical; [29]). The strength of such approaches, of course, is that they are based on the responses to actual odorants rather than to theoretical odorant constructs, and they can offer some predictive utility within localized regions of odor space. Efforts to combine receptor response data systematically with pharmacologically sound principles of ligand interaction is a critical but challenging process [30, 31].

We here present a general physicochemical olfactory sampling framework based on established pharmacological laws in which arbitrary combinations of odorant ligands and receptors can be constructed and their individual and collective effects on the resulting odor representation studied. Receptors occupy regions of a physicochemical quality space, or *Q-space*; this space maps the strength and efficacy of ligand-receptor interactions and enables recognition of the physical similarities among chemical species that are reflected in overlapping receptor interactions. Critically, odor perceptual similarities depend not only on their chemical properties, but also on the particular species-specific receptor complement deployed by a given animal. The axes of Q-space are not necessarily associated with any particular physicochemical property; they can be considered as fully arbitrary for theoretical work such as that described herein. Alternatively, if direct correspondence with real-world odorants is desired, Q-space axes can be defined as principal components derived from some similarity mapping process (e.g., [22–24, 26]), or as arbitrarily-defined molecular descriptors that may or may not correlate with any aspect of perceptual similarity. In either case, the dimensionality of Q-space and the sampling distributions of the odorant receptors deployed therein would be adapted to the datasets and questions of interest.

The interaction of odor ligands with these receptors is defined in the model by standard pharmacological laws. Odors may comprise one or many molecular types in characteristic concentration ratios; additionally, each component molecule may present multiple ligands to the complement of receptors expressed by a given animal. Individual odor ligands exhibit receptor-specific affinities and efficacies; that is, they may bind strongly or weakly to a given receptor, and can act as strong agonists, weak agonists, partial agonists, or antagonists [31–44]. Ligands interacting with common receptors compete with one another for dwell time; these competitive interactions appropriately simulate the degeneracy that fundamentally defines the capacities and limitations of odorant sampling. The outcome of these competing ligand-receptor interactions is expressed as a pattern of receptor activation levels, thereafter mapped to glomerular presynaptic activation levels based on the convergence of sensory neuron axons. The metric of greatest interest is the *mean discrimination sensitivity* 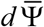, a measure of how effectively an olfactory system at this level is able to recognize a small change in the physicochemical quality of a stimulus.

This model presents several significant outcomes, both expected and surprising. First, adding additional receptors reliably improves the system’s discrimination sensitivity. Second, in contrast, adding additional ligands to an odorscene initially can improve discrimination sensitivity, but eventually will reduce it as the number of ligands increases. At very high ligand numbers, this process underlies and explains the phenomenon termed “olfactory white”, a phenomenon in which odorant mixtures with very large numbers of components at comparable intensities become increasingly perceptually similar to one another [27, 45]. Third, the presence of antagonistic ligand-receptor interactions generated several clear benefits to sensory system performance, producing higher absolute discrimination sensitivities and increasing the numbers of competing ligands that can be present before discrimination sensitivity began to be impaired. Finally, the model correctly reflects and explains the reduction in odor discrimination sensitivity exhibited by *Kir2.1* transgenic mice, in which the specificity of glomerular targeting by primary olfactory neurons is modestly disrupted.

## Methods

### Model overview

Simulations were designed in Python 2.7.13 as fully object-based models. A physicochemical metric space of odor quality (Q-space) was defined, into which odor ligands and the chemoreceptive fields of olfactory receptors were deployed. Realistic odorants comprised lists of odor ligands at characteristic concentration ratios. Ligands interacted with receptors according to standard pharmacological binding and activation laws and competitive interactions. Receptors were activated according to the outcome of their interactions with multiple ligands of varying affinities and efficacies. Glomeruli receiving input from single receptor types (*S*R*T*, the mammalian wildtype condition for the main olfactory bulb) directly inherited the activation level of their associated receptor type, whereas glomeruli receiving input from multiple receptor types (MRT, a condition exhibited by *Kir2.1* transgenic mice; [46]) inherited a weighted average of their constituent receptors’ activation levels. The *discrimination sensitivity* – the theoretical capacity of a network to discriminate small changes in odor quality based on particular odorants of interest, analogous to a least-discriminable difference measure – was measured and constitutes the primary output of interest. Receptor occupancy and receptor activation levels also were routinely measured. In the experiments described in this paper, odorants were based on lists of randomly placed ligands drawn from a uniform distribution across Q-space, such that the findings can be considered general. (The species-specific allocation of receptors to particular regions of Q-space at the expense of others – a foundation of natural scenes adaptation in olfaction – is a major motivation for development of this model; however, we here omit consideration of such heterogeneous Q-spaces in favor of first establishing general principles). Binding kinetics were omitted from the model (i.e., considered to be fast); detailed kinetic models of ligand-receptor interactions in olfaction have been developed by Rospars and colleagues; e.g., [47–49]. The objects and stages of the present model are described in detail below.

### Physicochemical model objects

#### Q-space objects

We first defined a physicochemical metric space object referred to as an odor “quality space”, or Q-space. Q-spaces were programmatically defined as *q*-dimensional metric spaces with arbitrary, finite ranges in each dimension. Within Q-space, proximity denotes unspecified similarities in the binding properties of chemical stimuli to receptors. Proximity also correlates locally with similarity in the naïve perceptual quality of odorants, but imperfectly so, as the distribution of receptors within Q-space strongly influences the perceived similarity of odorant inputs (as modeled by the transformation of the afferent representation into R-space; see *Discussion*). Moreover, other post-sampling factors such as learning and neuromodulation also affect perceptual quality in ways that are beyond the scope of this physicochemical model to address [12]. Q-space objects comprised lists of *q* tuples, each defining the range (minimum and maximum values) of one dimension of the space. The units of Q-space (q-units) are arbitrary, but the scale is shared among many model parameters such as the standard deviations of receptor Gaussians and the vector amplitude of *dN* (described below).

#### Receptor objects

Primary olfactory receptor (OR) objects were modeled as pairs of *q*-dimensional Gaussian distributions, both centered on the same point within a Q-space of dimension *q* (Figure 1). These two Gaussians defined the chemoreceptive field of the corresponding OR. Each of the pair of Gaussians exhibited independent, randomly-determined standard deviations in each dimension, and the profile of standard deviations across dimensions also differed between the two Gaussians of each pair. The first Gaussian defined in each OR object was an *affinity Gaussian* of dimensionality *q*, incorporating one *q*-dimensional coordinate defining the mean and a list of *q* independently-determined standard deviations. These objects also defined heuristic scalar values for the “strongest possible affinity” (at the mean/peak, in molar) and the “weakest possible affinity” (at the Gaussian asymptote). For simplicity in the present study, these values, defining the range of possible ligand-receptor dissociation constants *K_d_*, were kept constant across all ORs, from 1e-08 M at the peak to 1e+02 M at the asymptote, except when specified otherwise. Second, a separate *efficacy Gaussian* was centered at the same point as the affinity Gaussian; its peak denoted a full agonist (1, unitless) and the asymptote a full antagonist (limit = 0). Differences in the standard deviations between the affinity and efficacy Gaussians of an OR across *q* dimensions consequently provided points in Q-space corresponding to all forms of competitive ligand-receptor interactions with that OR (e.g., strong agonist, weak agonist, partial agonist, strong antagonist; Figure 1), and ensured that small changes in molecular features (as modeled by a 0.01 q-unit quality translation *dN* in *q* dimensions by a ligand point) could have either minor or major effects on a ligand’s binding properties to a given receptor.

**Figure 1.**
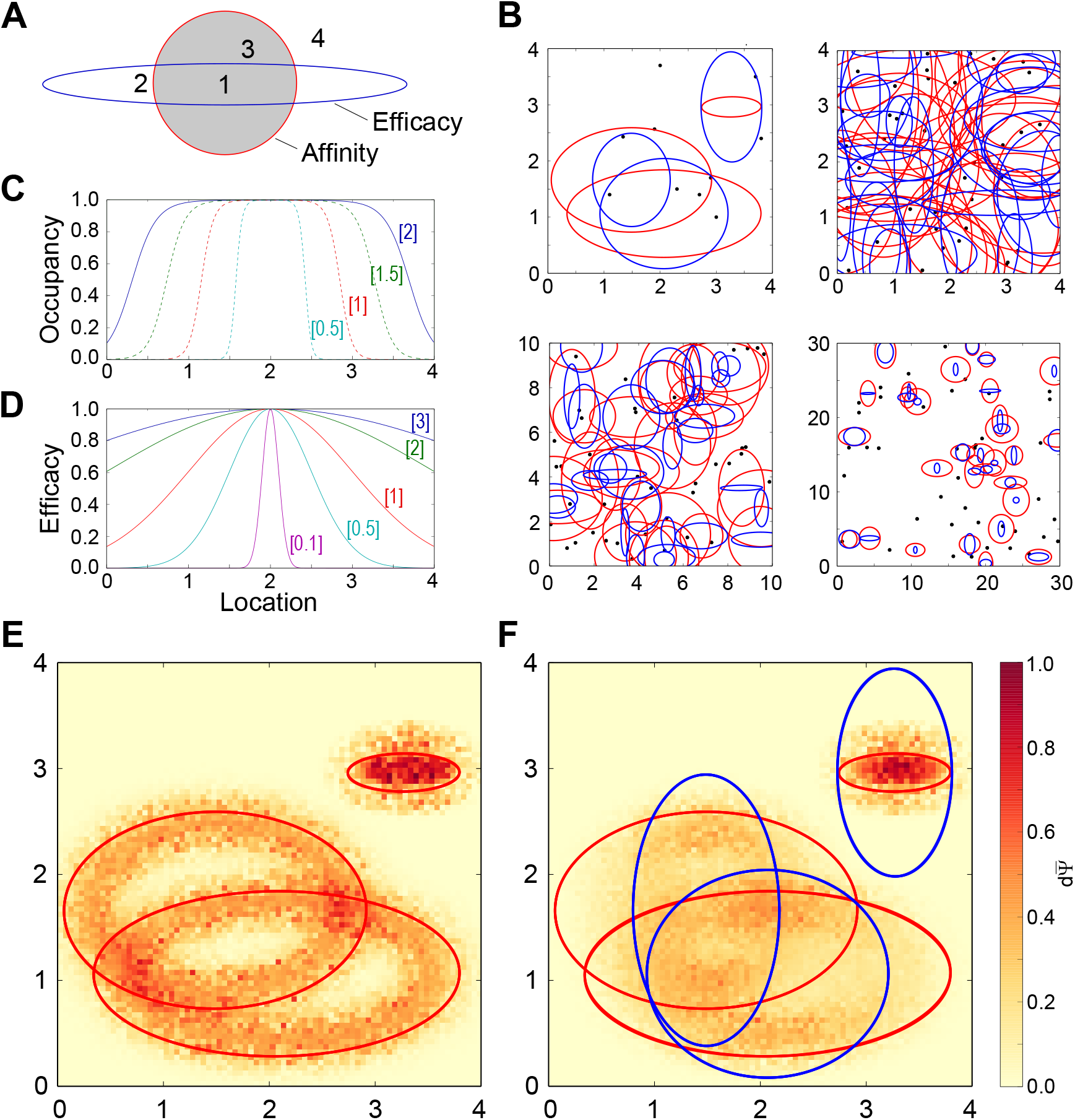
Model parameterization. **A.** Illustration of how the relationship between the affinity and efficacy Gaussians generates diverse ligand-receptor binding and activation properties. Black (affinity) and red (efficacy) ellipses denote the contour line marking two standard deviations (2.0 σ) from the Gaussian peak in a two-dimensional Q-space. *1*, Strong agonist (high affinity, high efficacy). *2*, Weak agonist (low affinity, high efficacy). *3*, Partial agonist or antagonist (high affinity, low efficacy). *4*, Weakly binding or insensitive (low affinity, low efficacy). **B.** Illustrations of receptor and ligand distributions randomly deployed from uniform distributions into two-dimensional Q-spaces of different sizes. Pairs of Gaussian contour lines (1.5 σ) centered on the same point denote the affinity (red) and efficacy (blue) Gaussians of individual receptors. Receptor standard deviations were drawn from uniform distributions of 0.5 to 1.5 q-units (affinity) and 0.05 to 1.0 q-units (efficacy; standard parameters). Black dots denote ligand points. *Top left*, A [4,4] Q-space containing four receptors and ten ligand points. *Top right*, A [4,4] Q-space containing 30 receptors and 40 ligand points. *Bottom left*, A [10,10] Q-space containing 30 receptors and 40 ligand points. *Bottom right*, A [30,30] Q-space containing 30 receptors and 40 ligand points. **C.** Effects of affinity Gaussian breadth on receptor occupancy. In a one-dimensional Q-space of range [4] containing a single receptor centered at location 2.0, with an affinity Gaussian exhibiting a standard deviation of 0.5, 1, 1.5, or 2 q-units and a fixed efficacy of unity, the receptor occupancy was calculated for 400 points distributed along the abscissa. Peak affinity was K_d_ = 10^-8^ M, asymptotic affinity was K_d_ = 10^+2^ M, and ligand concentration was 10^-5^ M (standard parameters). Occupancy fell off rapidly between about 0.7 and 1.0 standard deviations from the mean of the underlying affinity Gaussian. Bracketed numbers denote the standard deviation of the affinity Gaussian. **D.** Effects of the breadths of receptor efficacy Gaussians. Standard deviations of 0.1, 0.5, 1.0, 2.0, and 3.0 q-units are illustrated. Efficacy fell off most rapidly between roughly 0.5 and 2.0 standard deviations from the mean of the efficacy Gaussian. Bracketed numbers denote the standard deviation of the efficacy Gaussian. **E.** Heatmaps depicting 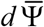 values per location in Q-space, with three receptors deployed and efficacy held at unity. Affinity Gaussians are depicted as red ellipses denoting 1.5 standard deviations (σ) on the affinity Gaussian function. Hot areas depicting higher discrimination capacities 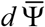 are clustered inside those ellipses, corresponding to the region with the steepest Gaussian slope (roughly 0.5 to 1.5 σ). Q-space locations served by two receptors exhibit correspondingly higher 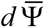 values at the intersections, and narrowly tuned receptors with correspondingly steeper Gaussians exhibit higher 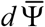 values at the cost of breadth. **F.** As E, with the addition of efficacy Gaussians (*blue ellipses*; 1.5 σ depicted). Scale bar on the right applies to both panels E and F.

This framework derives from the assumption that there exists a theoretical best ligand (strong affinity, efficacy of unity) for activating any given receptor. Any structural deviations from this theoretical optimum reduce affinity, efficacy, or both; accordingly, the peak affinity and peak efficacy coincide at the same point. That said, the underlying theoretical implication that agonists are, on average, slightly stronger than antagonists does not measurably affect model outcomes given the relative sparsity of ligand point deployment in Q-space. (Nevertheless, this simplifying assumption can be eliminated at the cost of adding a *q*-dimensional offset distance parameter between the peaks of the affinity and efficacy Gaussians to the receptor object). Deviations that reduce affinity but maintain efficacy create weak agonists; deviations that reduce efficacy but maintain affinity create strong partial agonists and antagonists. The number of independent ways in which a ligand can be so modified is reflected in the dimensionality of Q-space. The relative likelihoods of weak and strong agonists and antagonists, and the amplitudes of the quality transitions *dN* necessary to produce them, are arbitrary properties reflected in the relative standard deviations of the affinity and efficacy Gaussians across *q* dimensions.

Any number of ORs can be deployed into a given Q-space (Figure 1B). An *olfactory epithelium object* (aka *receptor complement*) was defined as a list of *κ* receptor objects, where *κ* = the number of receptors deployed in the model. A given olfactory epithelium object is considered characteristic of a particular animal species or population expressing a common complement of odorant receptors. Unless stated otherwise, simulations herein deploy 30 ORs into Q-space (i.e., *κ* = 30).

#### Ligand objects

Ligand objects consisted of a *q*-dimensional point coordinate (*ligand point*) deployed in a Q-space of dimensionality *q*, along with a concentration parameter in molar (M) specifying the concentration of the odorant molecule presenting the ligand. Mono- and multi-molecular odorants, odorant mixtures, and complex odorscenes all comprised simple lists of ligand objects (see *Discussion*). To model a single type of odorant molecule (monomolecular odorant) presenting multiple ligands, each ligand object associated with that odorant would have the same concentration value. To vary concentration ratios in binary odor mixtures within the model, the concentrations of the ligands comprising each of the two component odorants would be covaried. To model a natural odor source comprising multiple molecular species, the ligands corresponding to the different component species would be modeled as different concentrations in characteristic ratios. Increasing ligand concentration both increases the occupancy of well-tuned receptors, up to a maximum asymptotic level, and recruits additional, marginally-tuned receptors into the activated ensemble [50, 51]. In the present simulations, for simplicity, all odorant concentrations were held constant at 1e-05 M.

Small changes in the quality of a ligand were modeled as a 0.01 q-unit, randomly-oriented translation *dn* of that ligand point in *q* dimensions. In the present analyses, small changes in the quality of a natural odor or entire odorscene, denoted as *dN*, were modeled as small, randomly- and independently-oriented translations of *all* constituent ligand points. For simplicity, in the present simulations, all quality translations *dN* were of a constant magnitude, applied to all existing ligand points, and averaged over multiple independent random directions of translation.

#### Odorscene objects

Odor stimuli – whether comprising monomolecular odorants, more complex (multimolecular) odorants, or rich mixtures of odorants originating from multiple sources – were modeled as *odorscene* objects, which consisted of simple lists of ligand objects. Notably, odorants – even monomolecular odorants – can comprise multiple ligands; that is, different facets of an odorant molecule can separately interact with receptors. The interaction of a given odorant molecule with more than one receptor therefore arises by two means: (1) the presentation of multiple ligands by that molecule (separate ligand points in Q-space each interacting with receptor Gaussians) as well as (2) the interactions of any single ligand with more than one receptor (multiple receptor affinity Gaussians overlapping with a single ligand point in Q-space). Ligand object lists were additive in the sense that an odor mixture or complex scene would be defined by the simple concatenation of the ligand lists of its components. Because a given olfactory stimulus contains no *a priori* information regarding whether it comprises a monomolecular odorant, a complex odorant arising from a single source, or a complex mixture arising from multiple natural odor sources, the list of ligands presented as stimulus input to the model was termed an *odorscene* (Figure 1B). Specifically, the term is agnostic to whether that odorscene arises from a single or multiple sources, or to whether any subcomponent within it may have acquired associative meaning – it simply refers to “the complement of ligands interacting with the nasal epithelium”. A meaningful odor – such as orange or coffee – is referred to herein as a *source*.

#### Glomerulus objects

Glomerulus objects were defined to express the activation levels of olfactory bulb glomeruli when a given olfactory epithelium was stimulated with a given odorscene within a given Q-space. In the olfactory system, glomeruli are discrete regions of the olfactory bulb consisting largely of the axonal arbors of large, convergent populations of primary olfactory sensory neurons (OSNs) that normally each express the same type of OR. The number of glomeruli, therefore, corresponds to the number of ORs expressed by a given species of animal (though they may be duplicated; e.g., in mice, axons from the OSN population expressing a given OR usually converge onto two separate glomeruli located in the medial and lateral regions of each olfactory bulb; [52, 53]). In addition to activation levels, glomerulus objects consisted of a list of receptor objects and the proportional connectivity from each, as well as (X,Y) coordinates for the glomerulus’ position within the *glomerular array* on the olfactory bulb surface (used to organize any proximity-dependent computations of interest). In the present study, the number of glomeruli in the glomerular array was always the same as the number of receptors. In models of wildtype/SRT (single receptor type) mice, the list of receptor objects for each glomerulus object included only the primary receptor and a proportional connectivity of 1.0, whereas in models of MRT transgenic mice, in which multiple receptor types converge onto common glomeruli [46], eight secondary receptors in addition to the primary receptor contributed some level of input to each glomerulus.

### Ligand-receptor binding

Ligand binding to receptors was determined by the intersection of each ligand point with each affinity Gaussian within Q-space; specifically, the value of the affinity Gaussian at each such intersection defined a dissociation constant (*K_d_*) for that pairwise interaction. Hence, a given ligand could bind to any number of receptors with independently determined affinities, and a given receptor could interact with any number of different ligands. Receptor occupancy (*ϕ*) for each receptor then was calculated via standard pharmacological equations based on the concentration parameter of the ligand object. Specifically, for a single receptor/single ligand system:

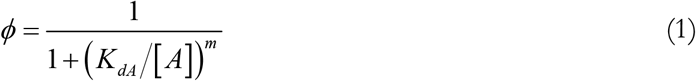

where [A] denotes the concentration of ligand A in molar and *m* is the Hill coefficient (cooperativity). The *partial occupancy* (ϕ*∂*) of any given receptor by any particular ligand in the presence of multiple competing ligands also was calculated using standard methods:

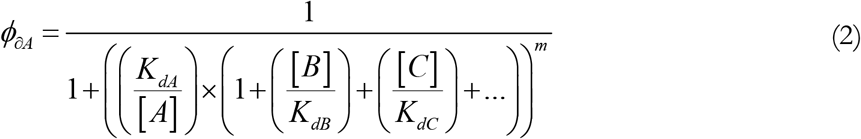

where A is the ligand for which partial occupancy is being calculated, and B and C are two other competing ligands; for each ligand species X, [X] denotes the concentration and *K_dX_* its dissociation constant with the receptor in question. This simplified equation assumes that all competing ligands exhibit the same cooperativity *m* when binding to the receptor, and neglects noncompetitive cobinding effects that may alter ligand affinities [54]. The total occupancy *ϕ* of a receptor in the presence of multiple ligands was calculated as the sum of all partial occupancies.

The value of the Hill coefficient *m* in isolated, deciliated OSNs has been estimated from 1.4 to over 4.4, based on physiological activation as opposed to binding [55]. Higher values result in sharper, narrower ligand-receptor activation curves. Populations of convergent OSNs, in contrast, reveal substantially lower functional Hill coefficients for collective activity at the glomerular level (H*ill equivalent*, [56]; i.e., broader dose-response curves [57–60]. While the model was designed to enable consideration of these parameters, in the present study the Hill coefficient was set to 1.0 (no cooperativity).

### Receptor activation

The contribution to receptor activation by each competing ligand was determined by ligand-receptor efficacy. Specifically, the efficacy of each ligand-receptor interaction was defined by the value of a receptor’s efficacy Gaussian at its intersection with the ligand point. For example, a ligand point that, owing to differences in the affinity and efficacy standard deviations, intersected with a high affinity value (low dissociation constant) and a high efficacy value would correspond to a strong agonist. Intersection with a low affinity and a high efficacy would denote a weak agonist, intersection with a moderate to high affinity and a moderate efficacy would constitute a partial agonist, whereas intersection with a high affinity and very low efficacy would correspond to a strong competitive antagonist. Importantly, the overall relative probabilities of agonistic and antagonistic interactions could be determined by varying the ranges of the distributions of values from which affinity and efficacy standard deviations were drawn (Figure 1A, C-D; Figure 2). For example, if affinity standard deviations were usually functionally narrower than efficacy standard deviations, then the prevalence of strongly antagonistic ligand-receptor interactions would be correspondingly low. The relationship of receptor activation levels to the location of ligand points in Q-space defines the tuning curves, aka chemoreceptive fields, of those olfactory receptors; i.e., OR tuning curves are a function of both the affinity and efficacy Gaussians.

**Figure 2.**
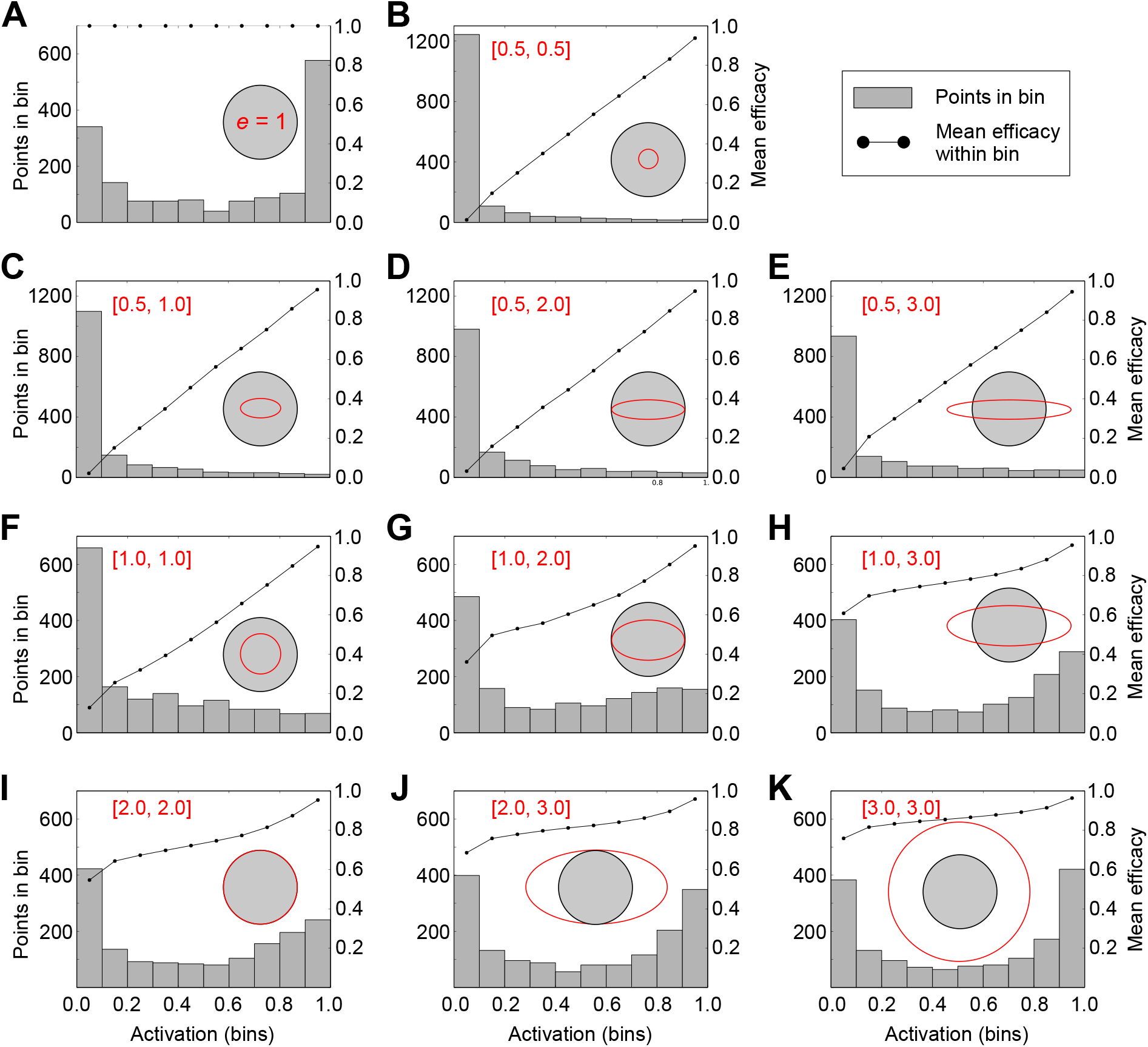
Depictions of the joint effects of receptor affinity and efficacy Gaussians on receptor activation. In each panel, a single receptor is deployed into the center of a two-dimensional [4,4] Q-space. The affinity Gaussian standard deviation (*grey shaded circle*) is uniformly 2.0 q-units in all cases, whereas the standard deviations of the efficacy Gaussian are systematically varied between 1.0, 2.0, and 3.0 q-units in each dimension (except in panel A, where efficacy is uniformly 1.0). A single ligand point was systematically varied across these Q-spaces in increments of 0.1 q-units (1600 points total per Q-space). The activation level of the receptor based on each of these 1600 ligands was calculated, and the results displayed as a histogram (*grey bars*; left ordinate). Broader efficacy Gaussians pull the distributions to the right, yielding higher probabilities of stronger receptor activation. Each of the ligand points also corresponded to an efficacy value; these values were averaged among all of the points within each of the ten bins of the histogram and plotted (*black curve*;, right ordinate). This reveals, for example, that the weakest activations in panel B are attributable to low efficacy (antagonist ligands), whereas comparably weak activations in panel K are associated with higher efficacy (agonist) interactions and hence their weakness must be attributable to low-affinity interactions (i.e., weak agonists).

### Discrimination sensitivity calculations

In order to measure the resolution of maximum odor discrimination afforded by a given receptor complement within a given environment, we developed a metric for least discriminable differences modeled after principles derived from the spectral overlap of cone sensitivities in the retina [61] and orientation tuning in primary visual cortex (V1; [62–64]). Specifically, the most sensitive odor quality discrimination thresholds occur where the *q*-dimensional OR tuning curves are steepest, rather than where they are most absolutely sensitive (Figure 1E-F). Multiple ORs that are sensitive to the same ligand further improve discrimination by contributing additional certainty (see *Results*).

We measured the *discrimination sensitivity* of a given receptor complement for a small change in the quality of a single ligand as the square root of the sum of squared deviations across all OR responses to a small, arbitrary translation *dN* of that ligand point in Q-space. Specifically, for a small *dN*, every OR generates a corresponding *dK_d_*, where *K_d_* is the amplitude of that receptor’s affinity Gaussian at its intersection with the ligand point (i.e., the ligand-receptor dissociation constant, in molar). That is, *dK_d_* represents the change in ligand-receptor binding affinity at one OR arising from the small transformation in quality *dN* of a single odor ligand.

Because the direction of a small translation vector *dN* in any one dimension of Q-space is arbitrary, and hence may be aligned with the steepest direction of the slope of a given receptor’s affinity Gaussian, or orthogonal to it (hence contributing zero *dK_d_* even if the affinity Gaussian is steep), or anywhere in between, a reliable estimate of the average discrimination sensitivity of a given OR complement for a given odorscene requires averaging across a number of random vectors *dN* from that odorscene. Additionally, as a given receptor complement will be better tuned to some odorscenes than to others, a reliable estimate of its discrimination sensitivity must be averaged across multiple randomized odorscenes. Figures depicting simulation results in this study are based on averages across 200 randomly determined odorscenes (each with *M* ligand points drawn from a uniform distribution across Q-space) with 10 random *dN* repeats per odorscene, sampled by a complement of 30 receptors randomly distributed within Q-space, unless stated otherwise.

Changes in ligand-receptor dissociation constants alone, however, do not fully determine discrimination sensitivity. First, discrimination across an odor quality difference *dN* is impaired when the odorant concentration is very low or high (saturating) compared to the dissociation constant, because small changes in the dissociation constant under these circumstances will have negligible effects on ligand-receptor binding and OR activation. Second, changes in ligand-receptor efficacy also can substantially influence the impact of small translations *dN* on OR activation. Third, odor sources generally comprise multiple ligands – potentially large numbers of ligands – and whereas multiple ligands may improve the discrimination of small quality changes by providing additional information, this will not always be the case. Additional ligands may saturate ORs, and, even in the absence of saturation, odor ligands that interact with the same receptor can interfere with one another; for example, increasing the affinities of an OR for both an agonist ligand and an antagonist ligand would generate a smaller-magnitude net change in receptor activation than would be the case for either ligand alone. To take all of these factors into account, we quantified the theoretical discrimination sensitivity of a receptor complement for an odorscene with *M* ligands, based on lists of length *M* of small randomly-oriented translations *dN*, as follows.

1. For each receptor object,

a. For each ligand point in the odorscene,

i. Apply the *dN* for that ligand point and determine the dissociation constant *K_d_* (intersection with the affinity Gaussian) for both states (i.e., *before* and *after dN* translation).
b. From these *K_d_* values and the corresponding ligand concentrations, calculate the following denominator term D for both states:

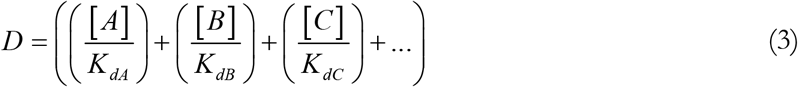

where D is the sum of [X]/K_dx_ for all ligands X.
c. For each ligand point in the odorscene,

i. Calculate the *partial occupancy* ϕ_∂,A_ of the current receptor by the current ligand A (using equation (2) substituted with the denominator term D from equation (3)) for both states:

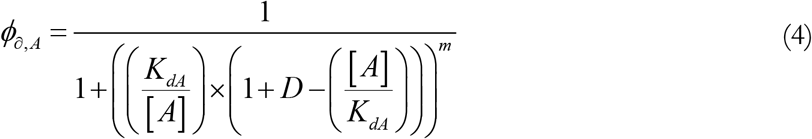
ii. Determine the efficacy *e* of the ligand-receptor interaction for both states (intersection with the efficacy Gaussian)
iii. Calculate the *partial activation* ε_∂,A_ of the current receptor R by the current ligand A, for both states:

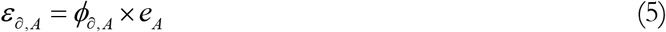

where *e*_A_ denotes the efficacy of the interaction of R and A. Importantly, while all ligands will contribute a nonnegative partial activation to each receptor, strong antagonists (for example) will reduce the partial occupancy of competing agonists, thereby reducing the level of receptor activation below that which would have been achieved in their absence.
d. Calculate the total receptor occupancy ϕ for the current receptor (i.e., the sum of partial occupancies for all ligand points), separately for both states.
e. Calculate the total receptor activation ε for the current receptor (sum of partial activations for all ligand points), separately for both states.
f. The change in total receptor activation dε in response to the quality translation *dN* is, therefore

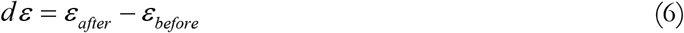 This value will be negative if the translation *dN* reduces net receptor activation.
g. The square of this value, *dε*^2^, is the positive, squared contribution of the current receptor to systemwide discrimination sensitivity.
2. The total discrimination sensitivity dΨ of a given receptor complement, for a given odorscene, in a given Q-space, based on a single set of ligand point translations *dN*, is therefore calculated as:

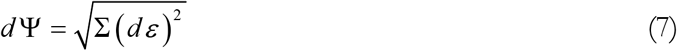 Note that *d*Ψ ≈ 0 in regions of Q-space in which no receptors are deployed (*approximately* zero because the long Gaussian tails of all receptors extend throughout the entirety of Q-space).
3. To characterize the discrimination sensitivity of a given receptor complement for a given statistical distribution of odorscenes, a mean discrimination sensitivity 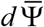 was calculated across 200 different M-ligand odorscenes drawn from a common distribution and 10 randomly determined translations *dN* of each odorscene. The mean discrimination sensitivity 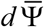 often is presented as a function of the number of ligands *M* per odorscene.

### The expected discrimination sensitivity 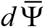

The mean discrimination sensitivity 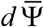 depends on receptor properties, numbers, and distributions as well as on odorscene statistics and environmental conditions. For example, nonuniform receptor deployments within Q-space would denote species-specific receptor complements tuned to favor the identification and discrimination of certain odor qualities at the expense of others, balancing improvements in discrimination sensitivity for favored types of odors against a reduced capacity to detect a wider range of odorants. In the present analysis, however, odor ligand and receptor deployments both were drawn from uniform distributions in order to establish the general principles by which olfactory discrimination sensitivity arises from basic pharmacological laws. Consequently, the *expected discrimination sensitivity* for a random odorant quality change *dN* is meaningfully equal to the mean discrimination sensitivity.

### The relationship between affinity and efficacy Gaussian functions

The model’s capacity to generate diverse ligand-receptor interactions (strong agonist, weak agonist, strong antagonist, etc.; Figure 1A-B) depends on selecting appropriately scaled standard deviations for the affinity and efficacy Gaussians in each of the *q* dimensions of Q-space. To determine these scales, we first calculated how the breadth of an affinity Gaussian corresponded to the breadth of a receptor occupancy function across Q-space. A single ligand point was deployed within a one-dimensional Q-space of range [4] expressing a single receptor with mean at location 2.0 and an affinity SD of 0.5, 1.0, 1.5, or 2.0 q-units. Efficacy was fixed at unity. The location of the ligand point was systematically varied from location 0 to 4 in increments of 0.01 q-units and the occupancy calculated at each location. In agreement with direct calculations (not shown), occupancy fell off rapidly between about 0.7 and 1.0 standard deviations from the mean of the affinity Gaussian (Figure 1C). In contrast, efficacy dropped off with distance as a direct function of the Gaussian, with most of the decline occurring between roughly 0.5 and 2.0 SDs from the mean (Figure 1D).

We then built a two-dimensional Q-space of range [4,4], deployed three receptors into it, varied the location of a single ligand point across the entire space in increments of 0.01 q-units, and calculated the discrimination capacity 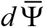 at each location. As expected, when efficacy was fixed at unity, higher values of 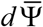 were calculated where the affinity Gaussian was steepest (roughly between 0.5 and 1.5 σ). The same relationship also has been observed for color discrimination performance in the human eye [65–67]. Redundant receptors increased the total 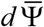 value at their intersections, and narrower chemoreceptive fields also generated correspondingly steeper Gaussian curves, thereby yielding higher 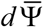 values (Figure 1E). When efficacy was permitted to vary, these functions further limited the regions exhibiting higher 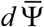 values (Figure 1F).

To demonstrate the effects of varying the standard deviations of affinity and efficacy Gaussians to generate diverse ligand-receptor interactions, we systematically built a set of two-dimensional Q-spaces of size [4,4], each containing one receptor comprised of an affinity Gaussian with a standard deviation of 2.0 in both dimensions (Figure 2; *grey shaded circles*) and one of a variety of efficacy Gaussians with standard deviations of 0.5, 1.0, 2.0, or 3.0 q-units in each dimension (Figure 2; *red ellipses*). The affinity SD of 2.0 was chosen because it fills the entire Q-space with occupancies above 0.1, with occupancy falling off most sharply between 1.4 and 2.0 q-units from the mean (Figure 1C, *solid curve*). We measured the activation ε of each receptor in response to stimulation with a single ligand point, the location of which was varied systematically across the entire two-dimensional Q-space in increments of 0.1 q-units; accordingly, 1600 activation values were plotted for each panel. For each receptor, the activation levels evoked by each of these ligand point locations were plotted together as a histogram (Figure 2; *histograms*). Notably, particularly when modeling stimulation with reliable agonists (e.g., when efficacy = 1.0; Figure 2A, or when the efficacy Gaussian was relatively broad when compared with the affinity Gaussian; e.g., Figures 2H, J, K), the distribution of receptor activation levels was noticeably bimodal. This straightforward result of interacting affinity and efficacy properties at the ligand-receptor interaction may contribute to a reported bimodality in mitral cell response properties that has been attributed to intraglomerular circuit operations [68].

The mean efficacy of all of the ligand-receptor interactions associated with each bin of each histogram then was calculated and plotted separately (Figure 2; *curves*). This demonstrates, for example, that the weakest receptor activations (leftmost histogram bin) in Figure 2K were primarily attributable to low-affinity interactions (weakly-binding agonists with high efficacy), whereas those in Figure 2B were largely attributable to low-efficacy interactions (antagonists). More generally, this figure shows that a balanced heterogeneity in affinity and efficacy standard deviations (e.g., Figure 2F-2H) generates diverse combinations of strong and weak agonist and antagonist interactions with receptors (Figure 1A). In subsequent simulations, except where otherwise specified, the affinity and efficacy standard deviations in each dimension were (separately) randomly drawn from uniformly distributed ranges rather than directly determined.

### Standard simulation parameters

We first characterized the physicochemical sampling properties of olfactory receptor complements with diverse properties. In each case, we modeled odor quality within a Q-space of a specified dimensionality *q* and range in each dimension. Thirty receptors were deployed into twodimensional Q-spaces unless stated otherwise. All affinity Gaussians were defined with a peak of 1e-08 M and an asymptote of 1e+02 M except where specified otherwise. The standard deviations of each affinity Gaussian in each dimension of Q-space were separately drawn from a uniform distribution ranging from 0.5 to 1.5 q-units, unless specified otherwise. Odorscenes ranged from 1 to 400 ligands (forming the abscissa of many plots); all ligands were presented at concentrations of 1e-05 M. The above parameters are considered *standard* and were utilized in all simulations depicted, except when specified otherwise.

Receptors’ efficacy Gaussians, when used, were centered on the same point in Q-space as the corresponding affinity Gaussian for that receptor. As described above, the ranges of standard deviations of the efficacy Gaussians were tuned against those of the affinity Gaussians in order to generate an appropriately diverse complement of ligand-receptor interactions. Based on this tuning, a standard range was determined from which the standard deviations for each dimension of receptor efficacy Gaussians were drawn. This standard range (0.05 to 1.0 q-units) was applied to all subsequent simulations except where specified otherwise. The peak of the efficacy Gaussian was always 1.0, and the asymptote always 0.0.

Unless stated otherwise, all plots were averaged across 200 odorscenes and 10 repeated *dN* translations per odorscene. In all changes in the quality of an odorscene performed herein, every ligand was translated by 0.01 q-units in a random direction. All receptors were deployed into locations in Q-space according to random sampling from a uniform distribution across the entire space; nonuniform sampling of Q-spaces was not performed.

### Animals

Mice carrying OMP-IRES-tTA:tetO-Kir2.1-IRES-tauLacZ alleles (Kir2.1 mice; [46, 69]) and wildtype controls were used as subjects. Kir2.1 mice were weaned on P21 and then maintained on a 20 mg/kg doxycycline-inclusive diet for at least six weeks prior to behavioral testing to induce the multiple receptor type (MRT) phenotype (Figure 8A). Littermate control animals are referred to as “single-receptor type” (SRT) mice. All animals were maintained in the Stowers Institute Lab Animal Services Facility on a 12:12 light cycle, and provided with food and water *ad libitum* except as described for two-alternative forced choice experiments. All behavioral experiments were carried out during the animals’ dark cycle under red or infrared illumination. All experimental protocols were approved by the Stowers Institute Institutional Animal Care and Use Committee (protocol 2019-102) and in compliance with the NIH Guide for the Care and Use of Laboratory Animals.

### Odorants

Monomolecular odorants were purchased from Sigma-Aldrich and diluted in mineral oil to the desired concentration as described below. Maple and lemon flavors were purchased from the Frontier Co-op (frontiercoop.com; cat #23081 and #23071) and were not diluted in the liquid phase. Odor delivery was controlled by an automated olfactometer with custom LabVIEW software (National Instruments) as described previously [26, 70]. All odorants presented were diluted tenfold in the gas phase by the olfactometer in addition to liquid-phase dilutions as noted. All mixtures were prepared in the liquid phase. Odorants are listed in Table 1.

**Table 1:**
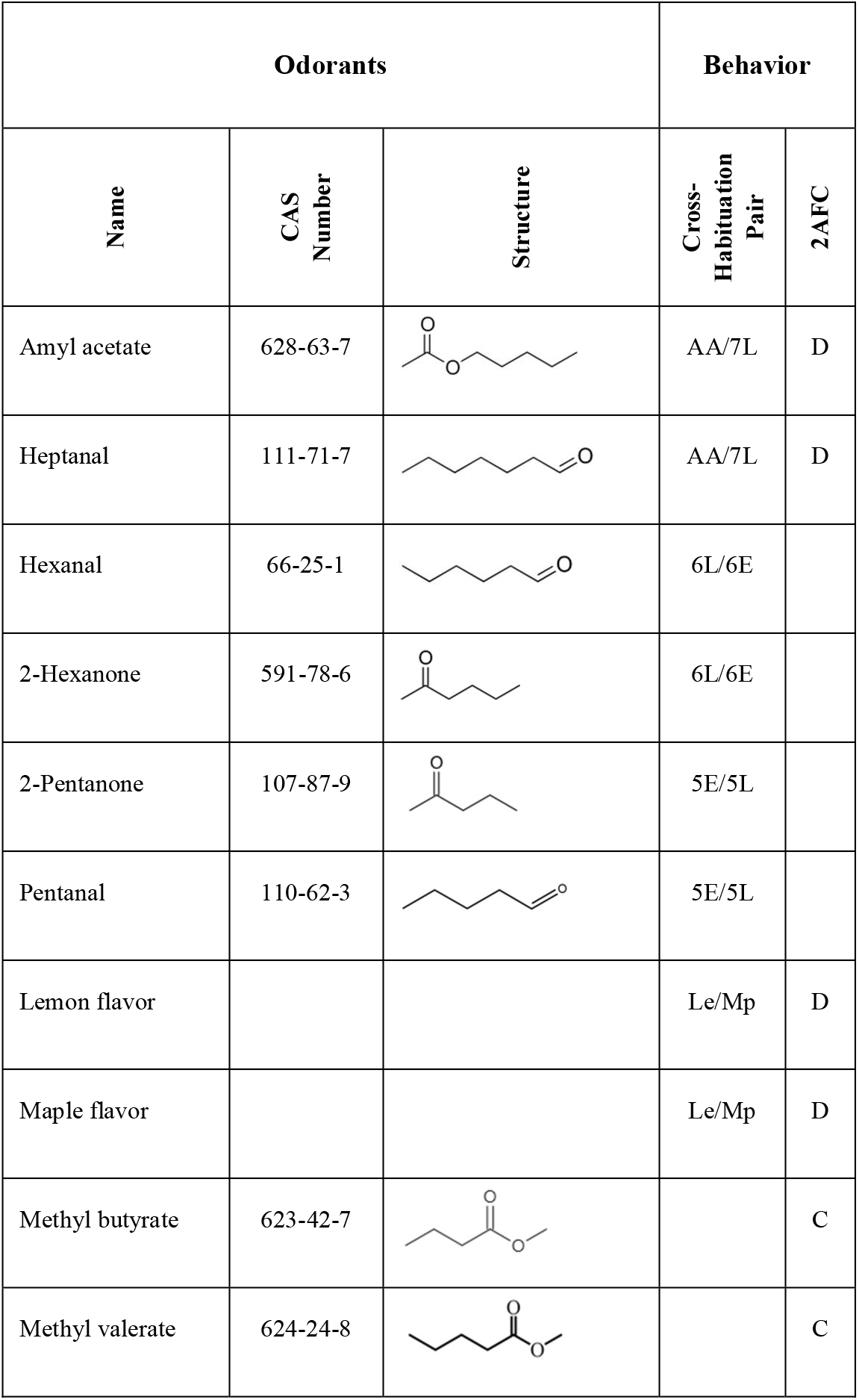
List of odorants. Odorants utilized in behavioral experiments along with their CAS numbers and chemical structures. The four odorant pairs used for cross-habituation studies (at 70:30 vs 30:70 mixture ratios; Figure 8B) are identified with descriptive labels. Odorant pairs used in two-alternative forced choice tests (2AFC; Figure 8C-D) are identified with their panel letter.

| Odorants |  |  | Behavior |  |
| --- | --- | --- | --- | --- |
| Name | CAS Number | Structure | Cross-Habituation Pair | 2AFC |
| Amyl acetate | 628-63-7 |  | AA/7L | D |
| Heptanal | 111-71-7 |  | AA/7L | D |
| Hexanal | 66-25-1 |  | 6L/6E |  |
| 2-Hexanone | 591-78-6 |  | 6L/6E |  |
| 2-Pentanone | 107-87-9 |  | 5E/5L |  |
| Pentanal | 110-62-3 |  | 5E/5L |  |
| Lemon flavor |  |  | Le/Mp | D |
| Maple flavor |  |  | Le/Mp | D |
| Methyl butyrate | 623-42-7 |  |  | C |
| Methyl valerate | 624-24-8 |  |  | C |

### Cross-habituation task

Cross-habituation tests were performed in 2-4 month old mice as previously described [70]. Each experimental group contained 6-14 animals. Each animal was tested with a total of 4 odors (2 pairs) in 2 separate experiments with at least one week between tests. Testing was performed in a 20 x 20 cm chamber to which the animals were first habituated for 30 minutes. Odorants were delivered by the olfactometer at 100 ml/min into a nose cone positioned on a side wall at 5 cm height. A vacuum tube connected to the opposite wall of the nose cone served to remove residual odors after odor delivery. Monomolecular odorants were diluted into mineral oil at 1:1000 (v/v), whereas lemon/maple flavors were not diluted in the liquid phase; for all odorants, a 10 ml/min air flow through the odorant-containing vials was mixed into 90 ml / min carrier air to yield a final nominal dilution of 10^-4^ saturated vapor (s.v.) for monomolecular odorants and 0.1 s.v. for lemon/maple flavors. Animals’ investigation of the odor source was registered by infrared beam breaking events and recorded by the same custom software that controlled the olfactometer.

In each trial, odorants were delivered for 1 minute followed by 4 minutes of carrier air. Mice first were habituated to the protocol by delivering eight trials of plain mineral oil, after which the first odorant of the pair (habituated odor) was presented for 5 trials and then the second odorant of the pair (test odor) was presented for 3-5 trials. During each trial, the duration of the animals’ investigation of the odor port was recorded. Repeated identical trials led to systematically reduced investigation (habituation), whereas perceived changes in the delivered odor restored the investigative response. Presentation of odorants perceived as similar to the habituated odor (i.e., not discriminated) evoke no increase in investigation time [1]. A normalized port investigation time (NPI) value was calculated for each trial by dividing the investigation time by the total baseline duration of odor port investigation during background delivery. The perceptual distance between the two odorants of a pair then was assessed as the difference in NPI (ΔNPI) between key trials.

Specifically, ΔNPI was calculated as the difference in investigation times between the fifth (last) presentation of the habituated odor (t_hab_) and the first presentation of the test odor (t_test_), expressed as a percentage over the average baseline exploration time during background air presentation.

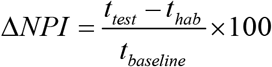

Values near zero therefore indicated a failure to discriminate between the paired odorants.

### Two-alternative forced choice task (2AFC)

Separate cohorts of 2-4 month old mice were used for this paradigm, in which mice are motivated to perform difficult odorant discriminations. In all cases, mice were water restricted (1.5 ml of water per day for one week) prior to training. Food was available *ad libitum*. Testing was performed in a 20 x 20 cm chamber with three nose cones positioned at 5 cm height. Odorants were delivered by the olfactometer at 100 ml/min into the central nose cone as described above, whereas the flanking nose cones were used to deliver water rewards. Mice were conditioned to nose poke into the central port, whereupon odorant delivery was triggered. Two odorants were presented in this fashion in a pseudo-random sequence, and water reward was contingent upon the animal correctly associating the odor with the appropriate water port. If the animal chose the correct port, 0.05 ml water was released; otherwise, no water was given. Animals were trained to criterion in the basic task (80% correct), before proceeding to the experimental paradigm.

The odorant discrimination success rate (SR) was calculated as

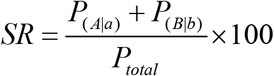

where *P*_(A|a)_, *P*_(B|b)_, and Ptotal were the number of pokes into water port *A* upon delivery of odorant *a*, the number of pokes into water port *B* upon presentation of odorant *b*, and the total number of nose pokes into the odor port, respectively. A success rate of 50% denotes chance performance.

In the first 2AFC experiment, 11 MRT and 18 wildtype mice were trained to discriminate the structurally and perceptually similar odorants methyl valerate and methyl butyrate at 1% saturated vapor (10^-2^ s.v.). Upon reaching criterion, discrimination performance at the testing concentration was assessed, after which the concentrations of both A and B were systematically reduced for subsequent testing at three lower concentrations (10^-3^, 10^-4^, and 10^-5^ dilutions). Liquid phase dilutions into mineral oil were 10% v/v for training and 1%, 0.1%, and 0.01% v/v for testing, each followed by an additional 10% dilution in the gas phase (see *Odorants* above).

In a second set of 2AFC experiments, mice were trained to discriminate pairs of odorants until achieving criterion (80%), after which they were systematically tested on their capacity to discriminate increasingly similar mixtures of the odor pair. For example, after being trained to discriminate amyl acetate from heptanal, 8 MRT and 8 wildtype mice then were tested on their ability to discriminate a mixture of 90% amyl acetate / 10% heptanal from a mixture of 10% amyl acetate / 90% heptanal, then 80:20 vs 20:80, and so on until reaching 50:50 vs 50:50, on which they performed at chance. All odorants/mixtures were presented at 10^-4^ dilution (1000x liquid phase dilution plus 10x gas phase dilution). This experiment was repeated in a separate cohort of 10 MRT and 10 wildtype mice using natural lemon and maple flavors, which were diluted only in the gas phase (10^-1^ overall dilution).

## Results

### Effect of ligand number and receptor density on odor discriminability with fixed full efficacy

We examined the relationship between the number of competing odorant ligands in an odorscene and the capacity of an olfactory system to discriminate small quality changes in that odorscene. To also assess the effects of *receptor density* (i.e., the extent of receptive field overlap in Q-space) without changing the absolute number of receptors, we constructed two-dimensional Q-spaces of three different sizes and deployed 30 receptors into each; i.e., the smaller Q-spaces exhibited higher receptor densities. Receptors were deployed randomly, with Gaussian peak locations drawn from a uniform distribution across Q-space. The standard deviations in each dimension of the affinity Gaussians were drawn from a uniform distribution ranging from 0.5 to 1.5 q-units. Efficacy was fixed at 1.0 (i.e., there was no efficacy Gaussian). We plotted the mean discrimination sensitivity 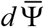 and mean receptor occupancy 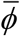 in each of these three Q-spaces as functions of the number of ligand points in the odorscene (Figure 3A).

**Figure 3.**
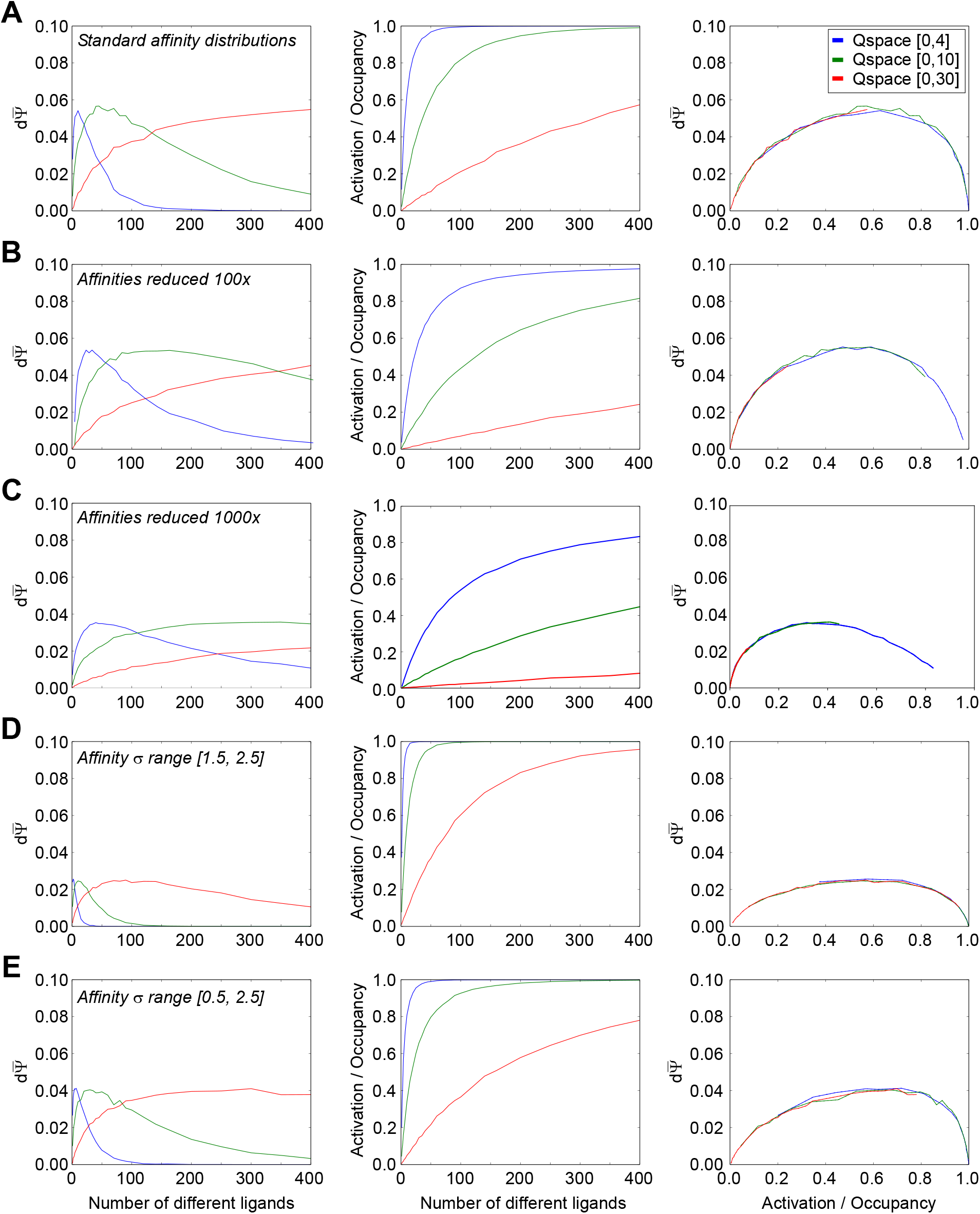
Effects of ligand and receptor densities on receptor activation/occupancy and discrimination capacity, while holding efficacy constant at unity. Because efficacy was held at unity, receptor activation equals receptor occupancy. Thirty receptors were deployed into each of three different Q-spaces in order to vary receptor density without changing the absolute number of receptors. **A.** Using the standard distribution of affinity Gaussians (σ = [0.5, 1.5]) and an efficacy of unity, the mean discrimination capacity (*left panel*) and receptor occupancy (*center panel*) were measured as functions of the number of ligands deployed in the space; the two functions also were plotted against one another (*right panel*). **B.** As panel A, but with affinities uniformly reduced by two orders of magnitude (Gaussian peak affinity K_d_ = 10^-6^ M, asymptotic affinity K_d_ = 10^+4^ M). **C.** As A, but with affinities reduced by three orders of magnitude (Gaussian peak affinity K_d_ = 10^-5^ M, asymptotic affinity K_d_ = 10^+5^ M). **D.** As panel A, but with the uniform distribution from which affinity Gaussian standard deviations were selected shifted to higher values, while retaining the same breadth of unity. **E.** As panel A, but with the uniform distribution from which affinity Gaussian standard deviations were selected doubled in breadth, incorporating the full range used across panels A and D. Extending the range of affinity standard deviations to higher values caused peak discrimination capacity to occur with lower numbers of ligands present (compare A to D, E), whereas excluding lower affinity Gaussian standard deviations (i.e., excluding receptors with sharper tuning curves) substantially reduced peak discrimination capacity (compare D to A, E).

Notably, discrimination sensitivity did not monotonically increase along with the number of ligands encountered in the environment, but rather peaked and then declined as the number of competing ligands continued to increase (Figure 3A, *left panel*). Indeed, at high densities of ligands, all delivered at similar concentrations, discrimination sensitivity was reduced to near zero; this reflects the phenomenon termed “olfactory white”, in which such mixtures become increasingly perceptually similar to one another [27, 45]. Reducing receptor density and receptive field overlap by increasing the size of the Q-space moved the 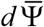 peak progressively towards higher numbers of ligands, indicating that the peak location was related to the number of ligands that substantively interacted with each receptor. One straightforward interpretation of this effect, all else being equal, is that higher numbers of interacting ligands result in higher receptor occupancies, which eventually approach saturation (Figure 3A, *middle panel*). That is, when the receptors are roughly half-occupied – i.e., near the steepest point in the underlying ligand-receptor binding curves – a small change in ligand quality *dN* generates a robust difference 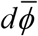. However, when those same receptors are nearly fully occupied by ligands, the same small change in *dN* will generate a near-zero 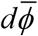, much the same as when the receptors are largely empty of ligands. This predicts that that the peak discrimination sensitivity 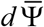 should coincide with receptor half-occupancy under these conditions. We tested this across different Q-space sizes, because the lower receptor densities presented by larger Q-spaces require a larger number of ligands to reach the half-occupancy point of any given receptor. Under these conditions, across a range of Q-space sizes, peak 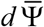 corresponded to approximately the number of ligands that delivered half-occupancy (Figure 3A, *right panel*), though this principle did not hold when efficacy was allowed to vary.

This relationship further suggested that lowering receptor occupancy values by reducing ligand-receptor affinities across the board, all else being equal, also should move 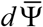 peaks to the right as they tracked the lower mean occupancies. To test this, we re-ran the simulations with the affinities of all receptors reduced by two (Figure 3B) or three (Figure 3C) orders of magnitude; that is, the peak of receptors’ affinity Gaussians now denoted dissociation constants of 1e-06 M or 1e-05 M, respectively, whereas the asymptotes now denoted 1e+04 M or 1e+05 M, respectively. With odor ligand concentrations still fixed at 1e-05 M, these modifications progressively reduced receptor occupancies (Figure 3B-C, *middle panels*), and demonstrated that 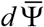 peaks continued to correspond to conditions in which receptors were half-occupied (Figure 3B; *right panels*). Notably, also, the peak discrimination sensitivity value was not strongly affected by the first 100x reduction in ligand-receptor affinity (compare Figure 3A and 3B) or the size of the Q-space, although the peak under these different conditions was reached under substantially different numbers of ligands. Further affinity reductions began to reduce peak discrimination sensitivity (Figure 3C).

We then measured the effect of increasing the mean breadths (standard deviations) of the affinity Gaussians in Q-space (cf. Figure 1C) while keeping constant the ranges of the distributions from which these standard deviations were drawn (i.e., the variability). Specifically, we drew the standard deviations in each dimension of Q-space uniformly from the range [1.5, 2.5] q-units (Figure 3D) rather than from the standard range [0.5, 1.5] depicted in Figure 3A. The range of affinities (dissociation constants) was maintained at standard values, consistent with Figure 3A; i.e., each receptor had a peak dissociation constant of 1e-08 M and an asymptote of 1e+02 M. This manipulation corresponded to increasing the breadths of olfactory receptors’ chemoreceptive fields, rendering them each responsive to a wider diversity of chemical ligands. Predictably, our simulations showed that this reduction in the selectivity of receptors for odor quality resulted in a substantial reduction in the aggregate discrimination sensitivity 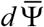 (compare ordinate values in Figure 3D, *left panel* to Figure 3A, *left panel*). This broadening of receptor chemoreceptive fields also shifted 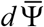 peaks to the left, corresponding to the increased levels of receptor occupancy generated by this broader selectivity (Figure 3D, *middle panel*). Peak 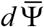 values again corresponded to conditions in which receptors were roughly half-occupied (Figure 3D, *right panel*).

Finally, we investigated the effect of broadening the range of the distributions from which affinity standard deviation values were drawn, to [0.5, 2.5] (Figure 3E), thereby encompassing the full ranges underlying the results from both Figures 3A and 3D. The increased diversity of receptive field breadths *per se* exhibited no specific effect; the occupancy and discrimination sensitivity functions of Figure 3E were intermediate between those of Figures 3A and 3D in both breadths and peak heights, as would be expected for a distribution with an intermediate mean.

Together, these results illustrate that discrimination sensitivity – for a specified number of receptors and in the absence of efficacy considerations or post-transduction neuronal computations – is determined by the average ligand-receptor occupancy functions of the deployed receptor complement. Specifically, discrimination sensitivity is highest under conditions in which mean receptor occupancy 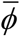 is near 0.5, corresponding to the steepest point in the sigmoidal ligand-receptor binding function.

### Effects of different receptor efficacy Gaussian functions

One critical limitation of the above analyses is that they omit the influence of ligand-receptor efficacy: the degree to which a ligand, once bound, will functionally activate the receptor. For any receptor, high-affinity, low-efficacy ligands (antagonists) can be found that bind strongly but do not open the receptor channel or activate its intracellular signaling cascade; also, many additional ligands exist that are of intermediate affinity and/or efficacy for any given receptor. We modeled the interaction between affinity and efficacy in Q-space by generating a second Gaussian for each receptor that determined the efficacy of any given ligand point when associated with that receptor (see *Methods*). Briefly, the standard deviations of a receptor’s efficacy Gaussian in each dimension were independent of each other and of the standard deviations of the affinity Gaussian; the resulting diversity in the relationships between the two Gaussians across Q-space generated regions for each receptor corresponding to the full range of possible competitive ligand-receptor interactions (Figures 1–2). Specifically, the relative breadths of the affinity and efficacy Gaussians determined the relative prevalence of agonistic versus antagonistic interactions in Q-space, and hence the relative likelihood of any given ligand being an agonist, weak agonist, partial agonist, or antagonist of a given receptor. We analyzed this relationship by applying efficacy Gaussians of different standard deviations (Figure 4A-C) or different randomized ranges of standard deviations (Figure 4D-F) to a set of receptors with standard affinity Gaussian heights and breadths drawn from the standard distribution (Figure 3A) to assess the impact of this set of variables.

**Figure 4.**
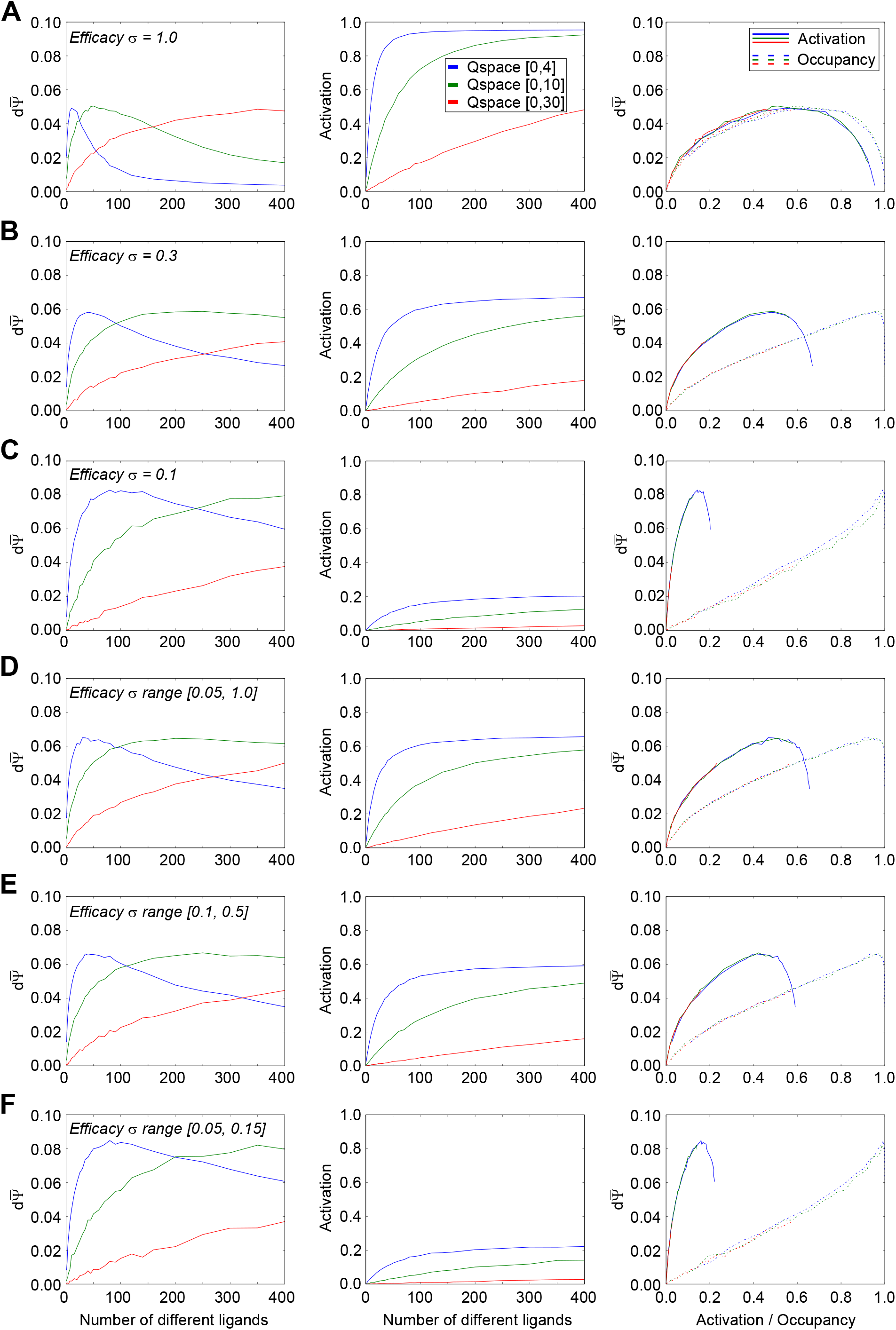
Effects of ligand-receptor efficacy distributions on receptor activation and discrimination capacity. All panels use the standard range of affinity Gaussian distributions (σ = [0.5, 1.5]; Figure 3A). Thirty receptors were deployed into each of three different Q-spaces. **A.** Discrimination capacity (*left panel*) and receptor activation (*center panel*) as functions of the number of ligands deployed when all efficacy Gaussians exhibit standard deviations of 1.0 q-units. The two functions also are plotted against one another (*right panel*). **B.** As panel A, except that all efficacy Gaussians are narrowed, each exhibiting standard deviations of 0.3 q-units. **C.** As panels A-B, except that all efficacy Gaussians are narrowed further, each exhibiting standard deviations of 0.1 q-units. **D.** As panels A-C, except that the standard deviations of efficacy Gaussians are drawn randomly from a uniform distribution with range [0.05, 1.0] q-units. This range is the standard parameter used to determine efficacy Gaussians unless specified otherwise. **E.** As panel D, except that the standard deviations of efficacy Gaussians are drawn from a narrower range of [0.1, 0.5] q-units. In concert, these results suggest that narrower efficacy Gaussians – that is, a higher prevalence of partial agonists and antagonists – increase the peak discrimination capacity despite exhibiting substantially weaker receptor activation levels, while also moving the discrimination peaks towards larger numbers of ligands. Additionally, as activation no longer directly tracks occupancy when efficacy varies, the relationship between activation, occupancy, and discrimination capacity became more complex. Peak discrimination capacity occurred at constant mean levels of receptor activation irrespective of receptor density *per se* (i.e., the size of the Q-space). Additionally, as efficacy Gaussians were narrowed (higher antagonist prevalence), the peak discrimination capacity occurred at correspondingly lower mean receptor activation levels. Finally, with a higher prevalence of agonists (broader efficacy Gaussians), higher occupancies reduced discrimination capacity well below its peak (A, *right panel*), whereas a higher prevalence of antagonists caused discrimination capacity to improve monotonically with higher receptor occupancy (C, *right panel*), presumably because quality shifts under those conditions are more likely to measurably alter net receptor activation levels.

For simplicity, we first looked at systems in which the standard deviation of the efficacy Gaussian was fixed, and also identical across both dimensions of the two-dimensional Q-space. Under standard affinity Gaussian parameters, efficacy Gaussians with standard deviations of roughly 3 q-units or higher (Figure 1D) presented receptor activation functions that were effectively identical to occupancy functions – i.e., ligands with appreciable dissociation constants would always have efficacy values of near-unity, hence the receptor activation function would closely track the occupancy function of Figure 3A, and the discrimination sensitivity functions therefore would closely resemble those of Figure 3A. Narrowing the standard deviation of a receptor’s efficacy Gaussian meant that a greater proportion of the ligand points exhibiting some nontrivial affinity for that receptor (i.e., that were close to the Gaussian peak location in Q-space) would be partial agonists or antagonists, as opposed to full (weak or strong) agonists.

As we reduced the standard deviations of the efficacy Gaussian, thereby increasing the probability of partial or antagonistic interactions, receptor activation levels (Figure 4A-C, *middle panels*) were correspondingly reduced (below their occupancies; the receptor *occupancy* function corresponding to all middle panels of Figure 4, which is independent of efficacy, is depicted in Figure 3A, *middle panel*). The number of ligands underlying peak discrimination sensitivity shifted to the right (Figure 4A-C, *left panels*), and no longer closely tracked either the point of 50% occupancy (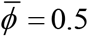; Figure 4A-C, *right panels, dotted curves*) or the point of 50% activation (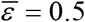; Figure 4A-C, *right panels, solid curves*). Specifically, as the probability of antagonistic interactions increased, the maximum activation levels of receptors at full occupancy were correspondingly reduced (*middle panels*), as were the absolute mean receptor activation levels at which peak discrimination sensitivity occurred (*right panels, solid curves*). The latter effect occurred for at least two reasons. First, 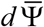 no longer declined to near-zero at near-complete occupancy, as was the case with full agonists only (Figure 3), because a quality translation *dN* now could result in agonists and antagonists replacing one another at the binding site, potentially altering receptor activation levels substantially without requiring a net change in receptor occupancy. Hence, the 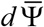 peak with respect to receptor occupancy moved to the right as the efficacy Gaussian narrowed (Figure 4B,C, *right panel, dotted curves*). Second, the increasing probability of antagonistic interactions imposed a correspondingly reduced maximum activation level on receptors; peak discrimination sensitivity reflected a balance between bound agonists and antagonists such that a quality translation *dN* was probabilistically optimized to impose the largest change in net activation per receptor. The 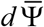 peak with respect to receptor activation therefore shifted to the left as the efficacy Gaussian narrowed (Figure 4B,C, *right panel, solid curves*).

Notably, we found that the highest absolute discrimination sensitivities were attained with narrower efficacy Gaussians – that is, ligand-receptor interactions with relatively high likelihoods of competitive antagonism (Figure 4C, *left panel*). Primary olfactory receptors, like any receptor, certainly have weak and antagonist ligands [32, 33, 37–39, 41–44], though the probabilities of each of these different ligand-receptor interactions for any given receptor complement clearly depend on the statistics of the odor environment in which it is deployed. In the present simulations, the maximum peak discrimination sensitivity was observed when the efficacy standard deviations were roughly 0.10 q-units (Figure 4C). Narrowing the efficacy Gaussian below a standard deviation of 0.05 reduced the peak 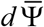 value, as antagonism began to dominate ligand-receptor interactions to the point of preventing appreciable receptor activation (not shown).

We then tested ranges from which to draw the standard deviations of efficacy Gaussians, in order to choose a standard range for subsequent simulations that would enable the system to exhibit fully diverse responses to odorscenes (Figure 4D-F). Specifically, we compared the full range of Gaussian breadths 0.05 – 1.0, from the value below which antagonism began to reduce performance to a value where, in the smallest Q-space used, efficacy would not decline below 0.1 (Figure 4D), to a more modest range in the center of this interesting region (Figure 4E), and to a narrower range encompassing only the values associated with the highest discrimination sensitivities (Figure 4F), among others (not shown). Based on these results, in order to make the fewest prior assumptions about the relative probabilities of agonistic and antagonistic interactions, for subsequent simulations we standardized on the full range of efficacy Gaussian standard deviations (0.05 – 1.0 q-units; Figure 4D). Individual standard deviations were drawn from a uniform distribution of values across this range.

The size of the Q-space (determining receptor density) had no effect on the relationships between 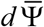 and receptor activation or occupancy (*right panels*); the curves depicted fully overlapped, though those associated with the larger Q-spaces did not always extend to the higher activation/occupancy values attained within the smaller Q-spaces given the maximum of 400 ligands tested. Of course, a given activation or occupancy would be attained at a larger absolute number of ligands in the larger Q-spaces, owing to the reduced overlap among receptors in those larger spaces. Finally, and importantly, these simulations make clear that the peak discrimination sensitivity exhibited by a given receptor complement does not generally occur either at 50% receptor activation or 50% receptor occupancy, as might be assumed a priori.

### Effect of ligand number and receptor density on odor discriminability with tuned affinity and efficacy ranges

Using the standard range selected for efficacy Gaussian standard deviations (0.05 – 1.0 q-units), we reproduced the simulations of Figure 3 incorporating the effects of these efficacy differences (Figure 5). The inclusion of diverse ligand-receptor efficacies modestly increased the peak discrimination sensitivity 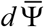 in all cases, and also deflected the 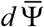 peaks toward higher numbers of ligands. This pattern held across reduced-affinity conditions (Figure 5B,C), and also persisted irrespective of increases in the mean breadths of the affinity Gaussians (Figure 5D) or increases in the heterogeneity of the breadths of the affinity Gaussians (Figure 5E).

**Figure 5.**
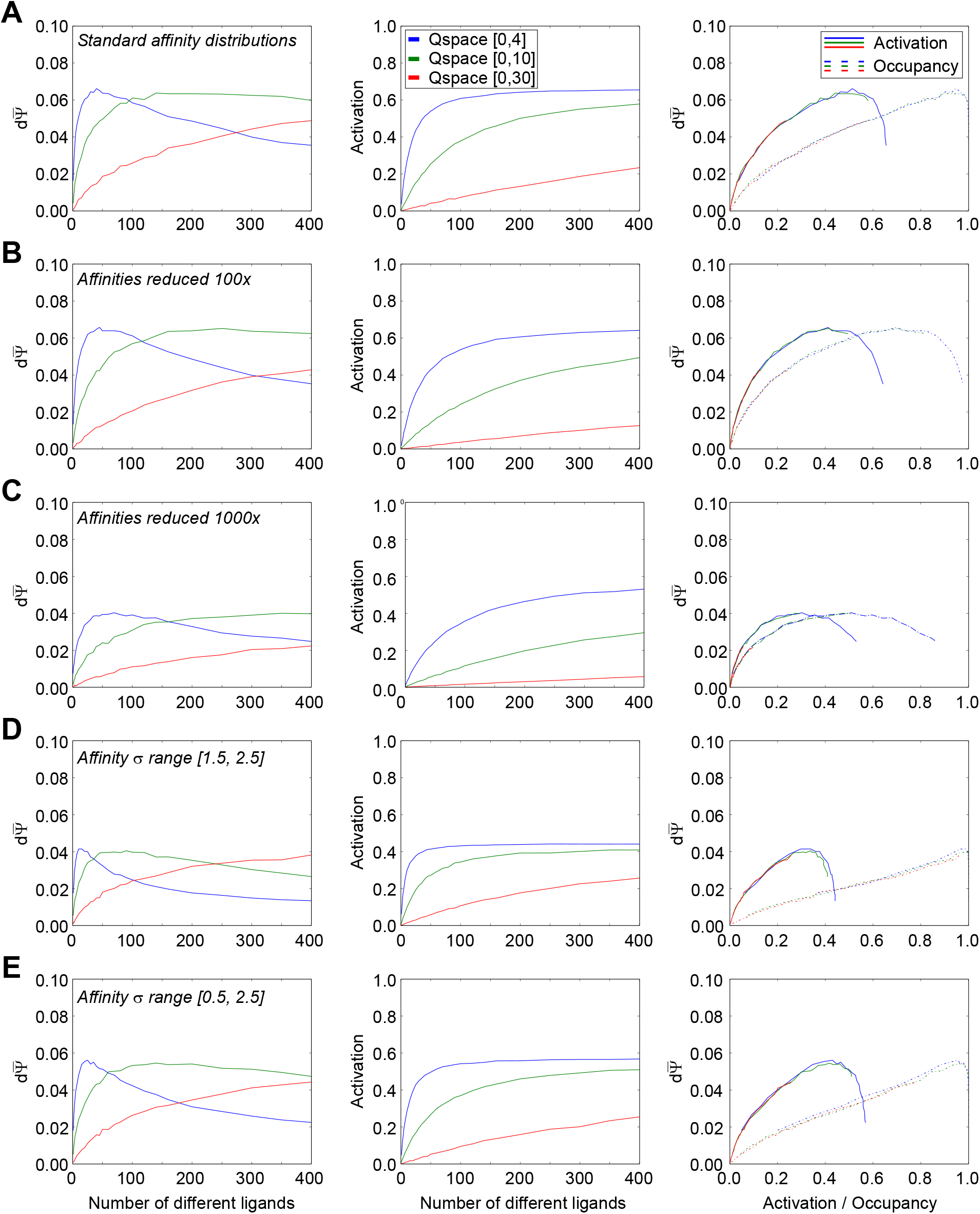
Effects of ligand and receptor densities on receptor activation/occupancy and discrimination capacity, while drawing receptor efficacy Gaussian standard distributions from the standard range of σ = [0.05, 1.0] q-units. Thirty receptors were deployed into each of three different Q-spaces. **A.** Using the standard distribution of affinity Gaussians (σ = [0.5, 1.5]), the mean discrimination capacity (*left panel*) and receptor occupancy (*center panel*) were measured as functions of the number of ligands deployed in the space; the two functions also were plotted against one another (*right panel*). **B.** As panel A, but with affinities uniformly reduced by two orders of magnitude (Gaussian peak affinity K_d_ = 10^-6^ M, asymptotic affinity K_d_ = 10^+4^ M). **C.** As A, but with affinities reduced by three orders of magnitude (Gaussian peak affinity K_d_ = 10^-5^ M, asymptotic affinity K_d_ = 10^+5^ M). **D.** As panel A, but with the uniform distribution from which affinity Gaussian standard deviations were selected shifted to higher values, while retaining the same breadth of unity. **E.** As panel A, but with the uniform distribution from which affinity Gaussian standard deviations were selected doubled in breadth, incorporating the full range used across panels A and D. Extending the range of affinity standard deviations to higher values caused peak discrimination capacity to occur with lower numbers of ligands present (compare A to D, E), whereas excluding lower affinity Gaussian standard deviations (i.e., excluding receptors with sharper tuning curves) reduced peak discrimination capacity (compare D to A, E). These effects are somewhat weaker than observed in Figure 3, in which efficacy was held constant at unity.

### Effects of increased numbers of receptors and glomeruli

In contrast to the nonmonotonic effects of increased numbers of odor ligands, deploying additional receptors into Q-space reliably and monotonically improved the discrimination sensitivity 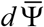 of the receptor complement, with negligible effects on the number of ligands or activation levels that were associated with peak discrimination sensitivity (Figure 6A-C; compare to Figure 5A, in which 30 receptors were deployed). That is, a greater metabolic investment to increase the number of odorant receptor types (along with the corresponding glomeruli and inter- and post-glomerular circuitry) will always pay dividends if the additional receptors interact with ethologically relevant odor ligands. Specifically, they will either improve discrimination sensitivity (as shown here, if deployed so as to overlap with the receptive fields of other receptors), increase the range of detectable odor ligands, or both. These results illustrate one of the reasons that olfactory systems exhibiting a greater diversity of odorant receptor types are superior, all else being equal.

**Figure 6.**
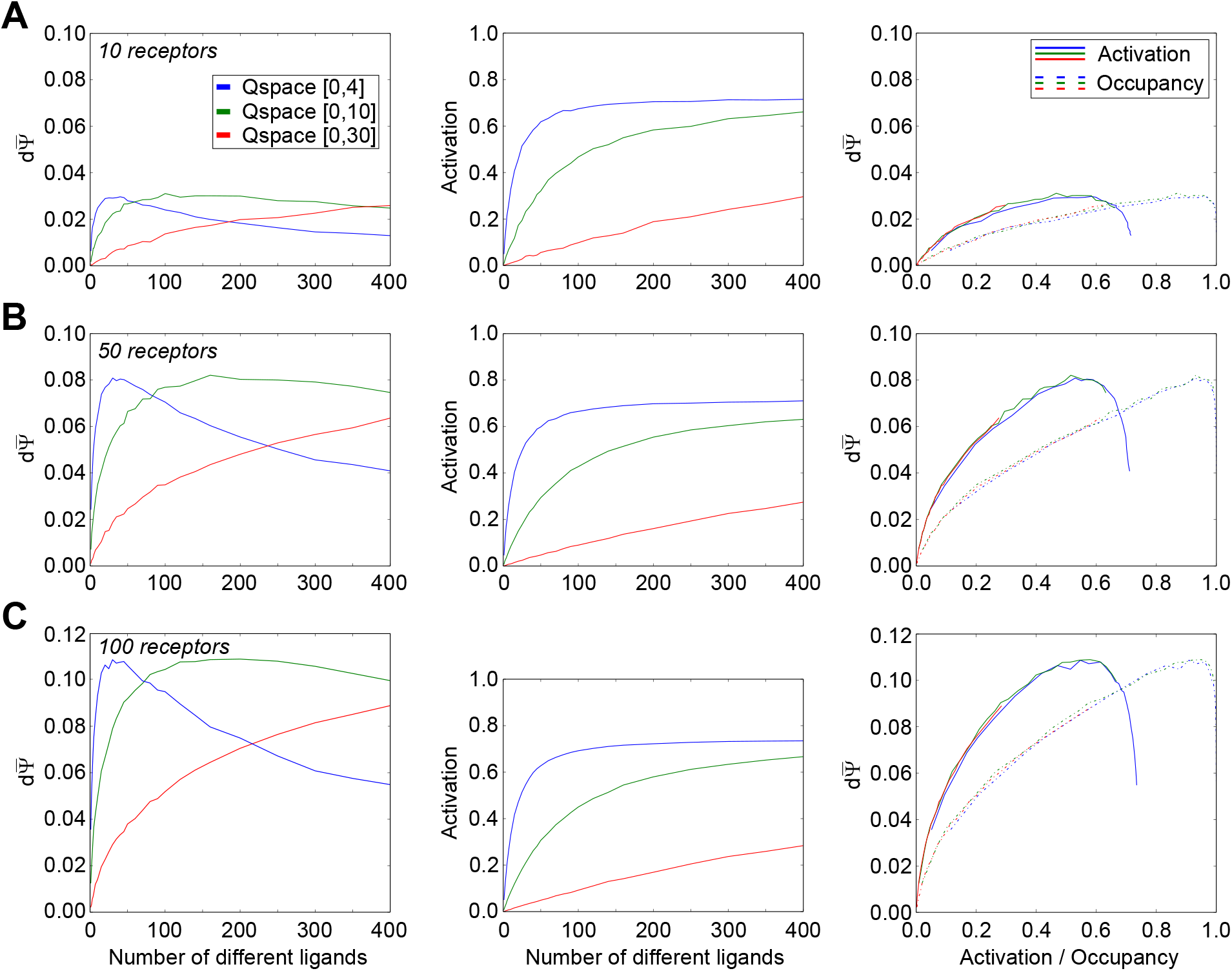
Effects of increasing the number of deployed receptors. In contrast to the nonmonotonic effects of increasing the number of ligands, increasing the number of receptor types deployed in a Q-space correspondingly improved the mean discrimination capacity in all cases. **A.** Receptor activation and discrimination capacity as a function of the number of ligands in a Q-space with ten receptors deployed. **B.** As panel A, but with 50 receptors deployed. **C.** As panels A-B, but with 100 receptors deployed. Compare also to Figure 5A, in which 30 receptors are deployed.

### Effects of higher Q-space dimensionalities

The dimensionality of Q-space is an important factor in modeling the complex effects arising from ligand-receptor interactions. Higher dimensionality rapidly increases the size of Q-spaces, reducing receptor overlap, but also increases the number of different relationships between affinity and efficacy Gaussian breadths that any one receptor can exhibit, and in principle increases the number of ways in which different receptors’ chemoreceptive fields can differ from one another. However, higher dimensions also require considerably more computational effort to converge on a clear mean outcome. Consequently, we asked whether, for the purposes of the simple questions asked herein, whether the basic response profiles of receptors deployed in higher-dimensional Q-spaces could be effectively replicated by receptors deployed in larger, two-dimensional Q-spaces.

We tested this inductively, by mapping discrimination sensitivity, activation, and occupancy functions in 2-, 3-, 4-, and 5-dimensional Q-spaces (spanning four q-units in each dimension; Figure 7A), and comparing them to those derived from larger, two-dimensional Q-spaces (Figure 7B). We found that many of the effects of higher-dimensional Q-spaces could be closely replicated by a correspondingly enlarged two-dimensional Q-space (compare curves in Figure 7A to 7B), echoing prior findings that representation accuracy by such metrics scales similarly irrespective of the dimensionality of the stimulus space [64]. The prominent exceptions to this general finding are the relationships between 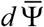 and the mean receptor occupancy and activation levels. While increasing the range of a Q-space does not systematically affect these relationships (Figure 7B, *right panel*), increasing Q-space dimensionality reduced peak 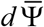 and substantially affected the mean activation levels at which the peak is obtained (Figure 7A, *right panel*). One reason for this is that increasing Q-space dimensionality, but not range, reduces heterogeneity in the outcomes of random ligand point transitions *dN*. That is, any translation *dN* in ligand quality generates an activation difference in each dimension which combine to generate the net activation difference *dε* of a given receptor. Importantly, the effects of a translation in one dimension may increase or oppose the effects of that translation in other dimensions. As dimensionality increases, the net variance of these superimposed effects will be reduced, therefore reducing the expected value of *dε* for a random translation *dN*. A second potentially contributing factor is that higher dimensions will statistically decorrelate the responses of proximal receptors to a single ligand translation *dN* – i.e., reducing the (already limited) extent to which a change in one receptor’s response could predict a change in the other’s response. The most appropriate dimensionality for Q-space is undefined, and indeed depends on details of the species-specific receptor complement and the list of odorants considered to comprise the olfactory environment; i.e., it is ultimately heuristic. For present purposes, we performed all of our simulations in two-dimensional Q-spaces except where otherwise specified.

**Figure 7.**
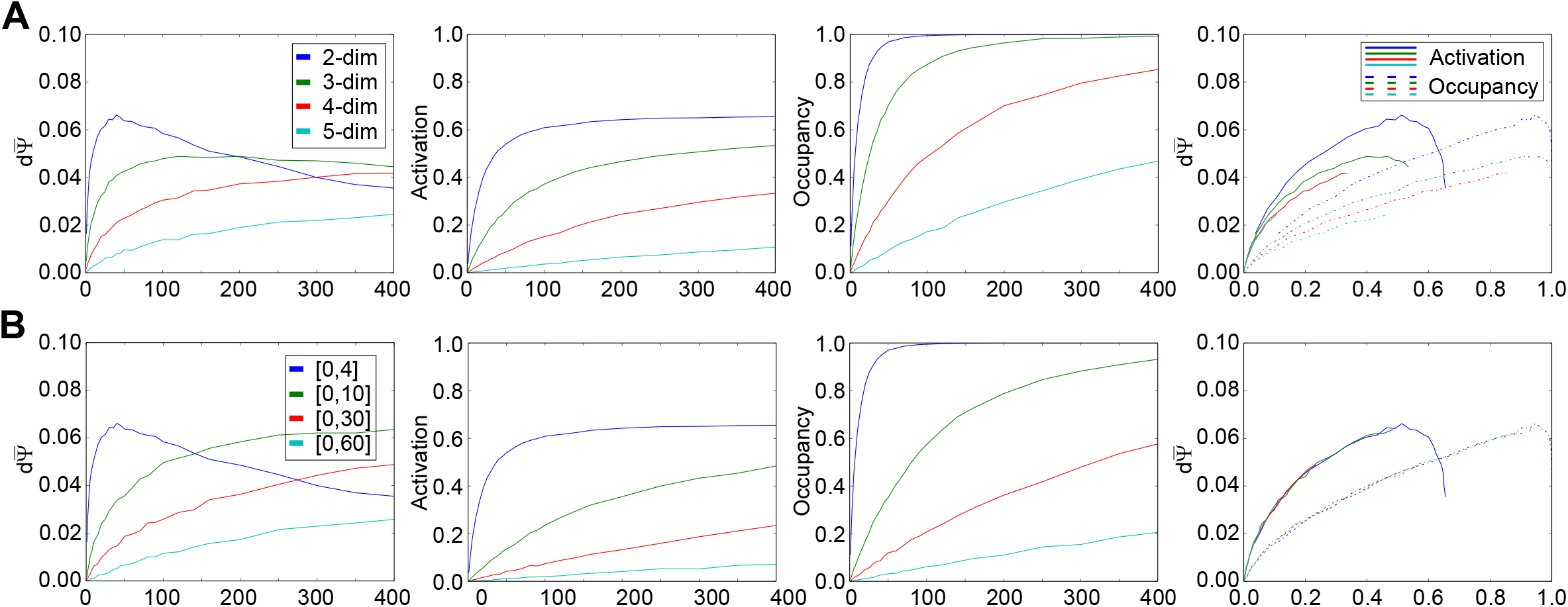
Effects of increasing Q-space dimensionality compared with increasing the size of a twodimensional Q-space. Receptor activation and receptor occupancy curves are separately depicted (*center panels*). **A.** Effects of increasing Q-space dimensionality. Each dimension spanned a range of four q-units, such that the three-dimensional Q-space had a range of [4,4,4], the four-dimensional Q-space was [4,4,4,4], etcetera. **B.** Effects of increasing the range of a two-dimensional Q-space. Most of the observed effects of higher dimensionality resembled the effects of simple increases in the size of the space, with subtle differences in curvature (compare *leftmost three panels*). However, Q-space dimensionality, but not range, affected the relationship between activation or occupancy and the discrimination capacity. Most significantly, increased dimensionality clearly reduced peak discrimination capacity, all else being equal, rather than simply moving the curve to the right so as to track receptor activation. This suggests that the larger number of factors contributing to net receptor activation changes based on any single ligand point’s transition *dN* tends to strengthen a regression to the mean effect, thereby reducing the likelihood of substantial changes in net receptor activation on which discrimination capacity depends.

### Effects of multi-receptor convergence onto glomeruli in MRT mice

In wildtype mice and other mammals, all primary olfactory sensory neurons (OSNs) expressing the same odorant receptor type converge axonally onto a common set of glomeruli in the olfactory bulb, and each glomerulus receives inputs only from that one class of receptor neurons. However, this rule is broken in transgenic Kir2.1 mice ([46]; see Methods), in which the receptor specificity of glomerular targeting is disrupted (Figure 8A); axons from OSNs expressing a given OR type project to several glomeruli (up to ~8) within a local region of the OB (although one glomerulus remains the primary target and some glomeruli may receive only a few fibers from that convergent OSN population). Conversely, a fraction of the axons converging on and constituting each glomerulus arise from OSNs expressing a number of different receptor types (*multiple receptor types* glomeruli; *MRT*).

**Figure 8.**
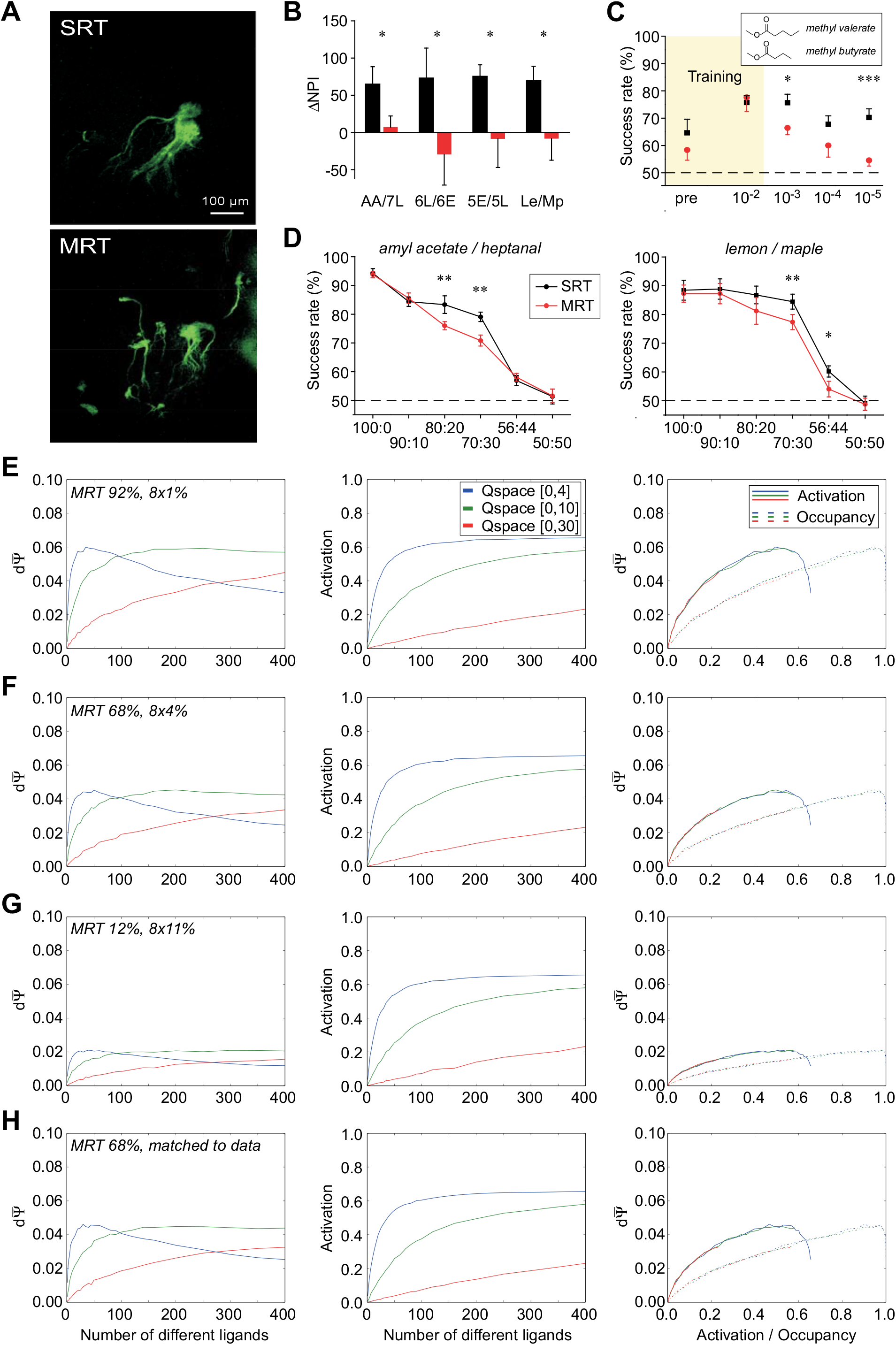
Effects of multi-receptor convergence onto common glomeruli. In contrast to wildtype (SRT) mice, in which each glomerulus is targeted only by one type of receptor, MRT transgenic mice exhibit glomeruli that are targeted by multiple receptor types. In the model, whereas each glomerulus still was predominantly targeted by a single corresponding receptor type, eight other receptor types also contributed smaller inputs. Mean occupancy, activation and discrimination capacity functions were calculated for glomeruli as weighted averages of those exhibited by their constituent receptors. **A.** Whole mount olfactory bulb images of green fluorescent protein signal from the axonal arbors of OSNs expressing the P2 receptor in control (SRT) and Kir2.1 (MRT) animals maintained on a doxycycline-inclusive diet for 8 months. Whereas P2 receptor axons converge onto a single glomerulus in SRT mice, this convergence is disrupted in MRT mice and multiple glomeruli are innervated to differing degrees; conversely, glomeruli are each targeted by multiple receptor types [46]. **B.** Cross-habituation between four variable-ratio odor mixture pairs (30:70 vs. 70:30) in SRT (black) and MRT (red) mice. Odor pairs are listed in Table 1 and were delivered at a dilution of 10^-4^, except for lemon/maple flavors (Le/Mp) which were delivered at 10^-1^ dilution (see Methods). MRT mice spontaneously discriminated each of the four odor pairs significantly more poorly than did SRT mice. **C.** Two-alternative forced choice odor discrimination performance when generalizing across concentrations. SRT (black) and MRT (red) mice were trained to discriminate between the structurally and perceptually similar odorants methyl valerate and methyl butyrate at a dilution of 10^-2^ (~1% saturated vapor; see Methods) and then tested on their discrimination performance at this concentration and at three lower concentrations. *Pre* denotes performance at the start of training. Dashed line denotes chance performance. **D.** Two-alternative forced choice odor discrimination performance as a function of odorant similarity, using variable ratios of binary odor mixtures to systematically vary similarity. Abscissa indicates the mixture ratio being compared to its complement; e.g., “90/10” denotes discrimination performance between a mixture of 90% amyl acetate / 10% heptanal and a mixture of 10% amyl acetate and 90% heptanal, whereas “70:30” denotes discrimination performance between a mixture of 70% amyl acetate / 30% heptanal and a mixture of 30% amyl acetate and 70% heptanal. Dashed line denotes chance performance. Significant post hoc pairwise comparisons between MRT and SRT mice in panels B-D are denoted with * for *p*<0.05, ** for *p*<0.01, and *** for *p*<0.001. Error bars denote SEM. Note that when presented with easier (less similar) odor pairs, MRT and SRT mice exhibit similar discrimination performance [74]. **E.** Occupancy, activation, and discrimination capacity effects in noses where each glomerulus received only 1% of its activation from each of the eight secondary receptors. **F.** Effects in noses where each glomerulus received 4% of its activity from each of the eight secondary receptors. **G.** Effects in noses where glomerular activation arose essentially equally from nine different receptor types. Greater overlap in receptor targeting of glomeruli yielded progressively inferior discrimination capacities (compare panels E, F, G). **H.** Effects in noses exhibiting a distribution of receptor contributions estimated from experimental data in MRT transgenic mice (*Kir2.1 mice*; [46]). Specifically, the primary receptor contributed a proportion of 0.68, and eight other receptors contributed proportions of 0.12, 0.08, 0.03, 0.03, 0.02, 0.02, 0.01, and 0.01, respectively. The model replicates the modest odor discrimination deficits observed in MRT mice compared with SRT wildtype animals (Figure 5A).

The specificity of OSN convergence onto individual glomeruli has long been hypothesized to contribute to odor discrimination capabilities [56, 71–73]. We asked whether a modest disruption of this convergence specificity on the scale demonstrated by Kir2.1 (MRT) mice would yield impairments in discrimination sensitivity when presented with tests rendered difficult owing to low concentrations or highly similar odorant mixtures; notably, Kir2.1 MRT mice exhibit no significant impairments in performance when presented with easier discrimination tasks [74]. First, we performed a cross-habituation assay between 30/70 and 70/30 ratio mixtures of four odor pairs (Table 1). Control (SRT) mice were able to distinguish the two mixtures reliably in all cases, whereas MRT mice performed significantly more poorly (two-way ANOVA with *genotype* and *odor pair* as factors; effect of genotype F(1,70) = 20.24, *p* < 0.0001; other effects not significant; Figure 8B). Using a two-alternative forced choice task, we then trained mice (to an 80% criterion) to discriminate between two structurally similar odorants presented at 10^-2^ s.v. (1%) concentration, and then tested their capacity to discriminate between these learned odorants at three lower concentrations. SRT mice remained able to distinguish the two test odorants at lower concentrations, whereas MRT mice performed significantly more poorly (two-way ANOVA with *genotype* and *concentration* as factors; effect of genotype F(1,138) = 17.71, *p* < 0.0001; effect of concentration F(2,138) = 4.24, *p* = 0.016, interaction not significant; Figure 8C). Finally, we used the same task to assess the mice’s ability to discriminate between highly similar odorant mixtures presented as a series of mixture ratios. A generalized linear model (binomial data, logistic regression) with two factors (*genotype, mixture ratio*) applied to data gathered from amyl acetate/heptanal mixtures (Figure 8D, *left panel*) indicated significant effects of genotype (χ^2^(1, n=16) = 6.38, *p* = 0.012), mixture ratio (χ^2^(3, n=16) = 247.81, *p* < 0.0001), and their interaction (χ^2^(3, n=16) = 12.50, *p* = 0.006). The same analysis applied to data gathered from lemon/maple flavor mixtures (Figure 8D, *right panel*) indicated significant effects of genotype (χ^2^(1, n=20) = 14.78, *p* = 0.0001) and mixture ratio (χ^2^(3, n=20) = 280.66, *p* < 0.0001); their interaction was not significant (χ^2^(3, n=20) = 1.58, *p* = 0.664). That is, for both odor pairs, MRT mice were impaired in their capacity to discriminate the more difficult mixture ratios compared to wildtype SRT mice (Figure 8D; asterisks denote significant post hoc pairwise comparisons).

To test whether a comparably modest disruption of this convergence specificity would yield an impairment in fine discrimination sensitivity in our physicochemical sampling model framework (i.e., a reduction in 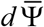), we constructed a range of OSN-glomerular projection profiles and measured their discrimination sensitivity functions. First, to directly examine the effect of glomerular sampling fields spreading out increasingly to incorporate multiple receptor types, we constructed MRT glomerulus objects (see *Methods*) in which the contribution of the primary odorant receptor was reduced from 100% (Figure 5A) to 92%, 68%, and 12%, with the eight glomeruli immediately surrounding the glomerulus of interest (in a 5×6 torus) each contributing an equal part of the remaining portion of receptor input (Figure 8E-G). All else being equal, broader sampling fields progressively reduced discrimination sensitivity in all cases, without affecting receptor occupancy or activation functions. We then constructed glomeruli with a “realistic” distribution of receptor inputs reflecting those observed in MRT mice (Figure 8H). Arrays of these MRT glomeruli yielded reduced discrimination sensitivities (impaired odor discrimination) compared with the SRT wildtype configuration (Figure 5A), reflecting the modest behavioral odor discrimination deficits observed in Kir 2.1 mice (Figure 8B-D). The performance of this distribution of receptor inputs was closely comparable to that of Figure 8F, suggesting that the relative contribution of the primary odorant receptor (68% in both cases) was the chief determinant of performance.

We then constructed a larger version of the “realistic” simulation, deploying 100 receptors into Q-space and mapping 100 corresponding olfactory bulb glomeruli onto a 10×10 torus. In separate simulations, for each glomerulus, we drew the minor receptor contributions either from the eight surrounding glomeruli (Figure 9A) or randomly from the entire glomerular field (Figure 9B). Drawing minor inputs to each glomerulus from random receptors yielded results no different than drawing them from the immediate physical surround of the primary glomerulus, as predicted given the absence of proximity-correlated chemotopy in the olfactory bulb [26, 75–77]. As expected, the peak discrimination sensitivities of this larger glomerular array were superior to that of the smaller 30-receptor deployments, but inferior to those of a 100-receptor deployment in a wildtype (SRT) configuration (Figure 6C). These results replicate, and potentially explain, the modest reduction in fine olfactory discrimination performance observed in MRT transgenic mice (Figure 8B-D).

**Figure 9.**
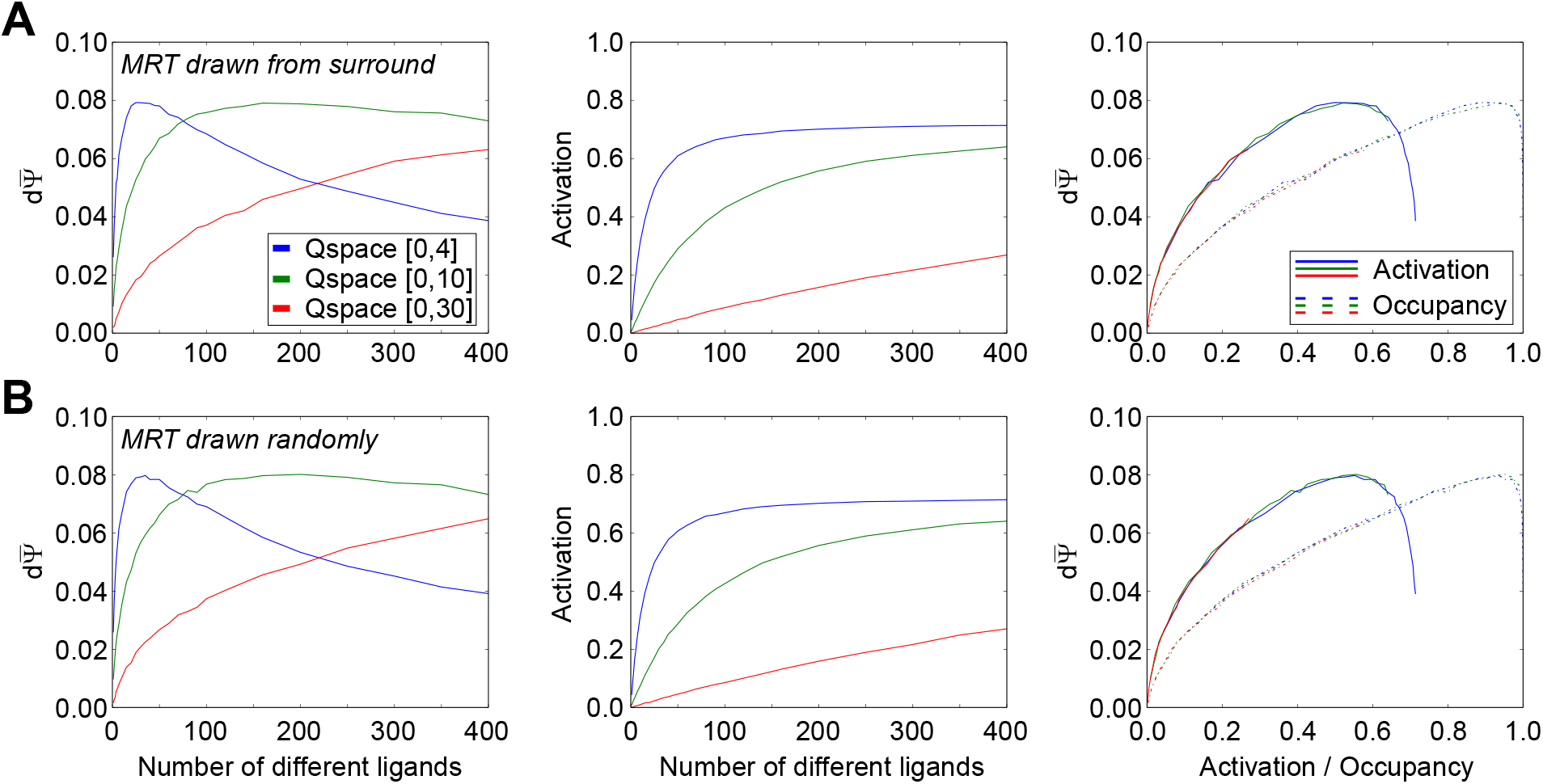
Absence of any effect of glomerular position on the MRT effect. To generate a 10×10 toroidal map of glomeruli, the number of receptors was increased to 100 for all plots. **A.** Restricting secondary glomerular inputs to those eight receptors primarily targeting immediately neighboring glomeruli yields patterns of neighborhood-based reciprocal relationships. **B.** Effects based on MRT glomeruli receiving their inputs from a random set of receptors. There were no discernable differences in discrimination capacity peaks or relationships between the two architectures. The relative contributions of the nine receptor inputs were based on experimental data as in Figure 8H.

## Discussion

This physicochemical sampling model simulates the pharmacological interactions between a species-specific complement of odorant receptors and an odorscene, which comprises one or more odor sources. Each odor source is based on one or more odorous molecules, each of which expresses some number of ligands via which it interacts with odorant receptors. This framework enables expression of the full complexity of odorant-receptor interactions based on the intersections of Gaussian receptor functions and lists of ligand points within a quality space of arbitrary range and dimensionality. The model enables assessment of the response of a given odorant receptor complement – i.e., the olfactory epithelium of a specific ‘nose’ – to a given odorscene, and how sensitive that response is to small changes in the odorscene. Figure 10 presents an example of this process of physicochemical sampling, illustrating the interactions of two multicomponent odors with a specific complement of odorant receptors, and how this interaction results in glomerular-layer odor representations that can be mapped into R-space. In the present simulations, ligands and receptors were deployed at randomized, uniform distributions across Q-spaces in order to examine the main effects of receptor properties and densities. Among other findings, the model demonstrates that the reduced glomerular targeting specificity exhibited by the *Kir2.1* (MRT) transgenic mouse model [46] would yield a modestly reduced sensitivity to fine odor discriminations, sufficing to explain that observed phenotype (Figure 8).

**Figure 10.**
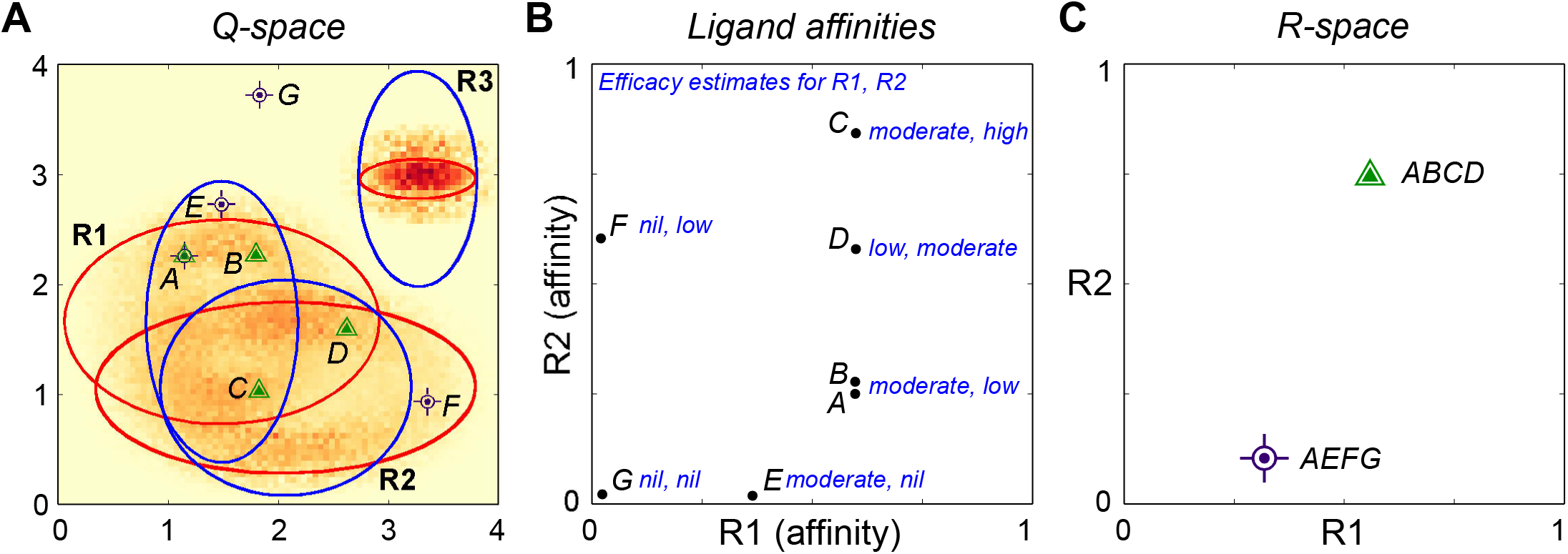
Illustration of physicochemical sampling and the transformation of odor representations from Q-space to R-space. **A.** A two-dimensional Q-space in which three receptors (Gaussian pairs R1, R2, R3) are deployed. Red ellipses denote the 1.5 σ contours of the affinity Gaussians, and blue those of the efficacy Gaussians, as in Figure 1. Two four-component source odors (i.e., comprising four ligand points) also are depicted: odor ABCD (*green triangles*) and odor AEFG (*violet bullseyes’*, note that the two odors share one ligand in common). For present purposes, all ligands are presented at the same moderately effective concentration. As shorthand descriptors, affinity and efficacy are here described as high or strong when ligand points are near the center of the respective ellipse, moderate when just inside the ellipse contour, weak or low when just outside the contour, and nil when distant from the ellipse. Ligand A has moderate affinity for R1 and activates it with moderate efficacy, so constitutes a reasonably effective R1 agonist. It binds only weakly to R2, and with low efficacy. Ligand B binds to R1 with the same affinity and efficacy as ligand A, and interacts very weakly with R2 (though marginally less weakly than A). Ligand C also binds to R1 with the same moderate affinity and efficacy as ligands A-B, but binds strongly and with high efficacy to R2. Ligand D again has the same moderate affinity to R1, but exhibits a much lower efficacy; it is essentially an antagonist of R1. At R2, ligand D binds moderately well and is an agonist of moderate efficacy. Ligand E binds more weakly to R1 than ligands A-D, but exhibits the same efficacy there as ligands A-C. It does not appreciably interact with R2. Ligand F does not interact appreciably with R1, but exhibits moderate binding and low efficacy to R2 – i.e., it is a partial agonist of R2 tending towards antagonism. Ligand G does not interact measurably with either R1 or R2 – that is, this particular complement of receptors is insensitive to ligand G. (Of course, a different animal species might express a receptor that interacts substantially with ligand G). None of the ligands interact measurably with receptor R3. The color of the heat map (see Figure 1F for scale) denotes a 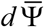 value for that ligand point – the expected contribution of that ligand (across all receptors) to the overall discrimination capacity. As the ligands of odor ABCD, combined, exhibit a greater sum of individual 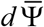 values than those of odor AEFG, it will be easier for the animal expressing these receptors to detect a small quality change to ABCD than it would to detect a similar quality change to AEFG. **B.** Plot of the affinities of each ligand for receptors R1 and R2, where 0 denotes no affinity and 1 denotes the strongest affinity (K_d_ = 10^-8^ M in most of the present simulations). Receptor R3 is neglected as it interacts with no ligands. Each plotted point also is labelled with that ligand’s efficacy at R1 and R2 (respectively). **C.** Depiction of the two source odors in the R-space defined by receptors R1 and R2 (R3 is again neglected). Odor ABCD (*green triangle*) activates R1 moderately, assuming that the three moderate agonists outweigh the effects of the single moderate antagonist, and activates R2 moderately strongly, because ligands C and D are both effective agonists whereas ligands A and B do not strongly interact with that receptor. Odor AEFG (*violet bullseye*) activates R1 relatively weakly, because A is a moderate agonist, E is a weak agonist, and F and G do not appreciably interact with the receptor. Among the ligands of odor AEFG, only ligand F interacts substantially with R2, as a low-efficacy partial agonist – consequently, R2 is minimally activated by that odor. The distance between these two odor representations in R-space denotes their physical similarity after sampling. Importantly, if these same two odors were sampled by an animal expressing two different receptors, this similarity relationship would be different. Moreover, the full representation of odors ABCD and AEFG in R-space does not actually comprise a single point, but a smooth manifold, based on experience and incorporating the natural quality variance of the odor source (discussed in [78]).

It is sometimes underappreciated that odorant-receptor interactions are simple ligand-receptor interactions that follow standard pharmacological laws. Whereas odorant receptors are sometimes characterized as “broadly tuned”, in supposed contrast to other seven-transmembrane receptors such as muscarinic cholinergic receptors, beta-adrenergic receptors, and metabotropic glutamate receptors, there is nothing special about odorant receptors save their diversity and their deployment in an unregulated external environment. Odorant receptors indeed respond to ranges of similar odorants, as well as to less-similar odorants that presumably exhibit some common ligand [30–33, 39], but this breadth is not a property of the receptors *per se* so much as it is a property of the statistics of their chemical environment. To wit, muscarinic cholinergic receptors also respond to several experimental agonists, and bind experimental antagonists, and there exist many more weak and partial agonists for this receptor that are not scientifically useful. However, the receptor is deemed a cholinergic receptor because it never encounters any of its ligands other than acetylcholine under natural circumstances. In contrast, odorant receptors are deployed into an unregulated chemical environment in which encounters with their diverse ligands are part of normal function. The present model framework is designed to quantify and explain such phenomena as the effects of larger numbers of ligands, different receptor complements, and the relative prevalence of antagonistic interactions on olfactory system function.

### Perceptual spaces

Quality space, or Q-space, provides a representation of physicochemical similarities among potential ligands. Critically, Q-space does not define odor perceptual similarity; that is a postsampling phenomenon that depends on the chemoreceptive fields exhibited by an animal’s receptor complement as well as on subsequent learning-dependent transformations in the brain. Rather, Q-space is defined by structural commonalities that underlie correlations in ligand-receptor binding likelihoods. For this reason, individual receptors are defined by a single pair of Gaussians localized within Q-space. If two molecular ligands are able to interact with a given receptor, then they are correspondingly close in Q-space, irrespective of how similar they might be in terms of some other arbitrary physicochemical metric. The large number of different ways in which molecular ligands are able to vary, and the diverse effects that any given change in a ligand might exert on any two different receptors, requires that Q-space be of an arbitrary dimensionality defined by the complexity of the odorscenes and the sets of odorant receptors to be investigated. Most simulations herein were performed in two-dimensional Q-spaces for ease of illustration, bolstered by results indicating that most outcomes in higher-dimensional Q-spaces also can be obtained in twodimensional Q-space (Figure 7). Interestingly, early efforts to map a physicochemical quality space based on perceptual similarities argued that most of the variance could be encompassed by two dimensions [22, 24], though the underestimation of dimensionality can be a common consequence of undersampling ligand diversity.

Odorscenes of arbitrary complexity comprising any number of independent sources are depicted as lists of ligand points within Q-space, each exhibiting a concentration. Notably, any combination of odor sources, irrespective of the complexity of each source or the number of sources present, can be decomposed into a simple list of ligands (with associated concentrations). Ligands that compete for the same receptors, whether arising from the same or different sources, will be adjudicated by standard pharmacological laws. This direct implementation also enables the straightforward modeling of corner cases. For example, monomolecular odorants with nonmonotonic doseresponse curves can be implemented as one high-affinity, high-efficacy ligand plus one somewhat lower-affinity, low-efficacy ligand; at higher concentrations, the latter competes effectively with the former to reduce the total receptor activation below its lower-concentration peak. Ultimately, these interactions between ligands and receptors generate a characteristic pattern of activity across the complement of receptors that comprises any given nose.

Receptor space, or R-space, is the sensory metric space into which the primary olfactory sensory response pattern is deployed [78]. Unlike Q-space, R-space has a defined dimensionality equal to the number of different receptor types expressed by the species in question (e.g., 1000-1200 in mice, ~400 in humans), such that any instantaneous pattern of response intensities among receptors, or glomeruli, can be depicted as a point in this unit space, within which a fully-activated receptor returns a coordinate of 1 in that dimension whereas an inactive receptor returns a coordinate of zero. This post-sampling sensory space is the outcome of the present model framework and the basis for subsequent neural computations that construct odor representations and perceptual similarity. R-space also is the basis for arguments that early olfactory computations are intrinsically high-dimensional [75, 76, 79–81]. Experimentally, the most direct (albeit partial) measurement of activity in R-space is the pattern of glomerular activation levels observable via optical imaging [26, 57–60], although the transformation from receptor activation to glomerular activation is nonlinear, including axonal convergence from different regions of the epithelium, feedback-based normalization [82], and other potential heterogeneities, some of which may underlie the broader dose-response curves observed via optical imaging at the collective glomerular level [4, 56]. In the present model, glomerular activation levels directly reflected receptor activation levels, except in simulations of transgenic MRT mice in which they reflected weighted averages of the multiple receptors contributing to their activation (Figure 8E-H).

### Ligand models vs. the theory of odotopes

The *ligands* that comprise odor sources in this model are subtly but distinctly different from *odotopes* following the definition of Cleland [83]. Briefly, the odotope there is defined as “the net effect of a given odorant molecule on a single type of odorant receptor”, divorced from a strict association with a particular binding ligand on that molecule. This definition offers clear theoretical benefits for incorporating quantitative biological data into non-pharmacologically based models. However, in the present circumstances, the goal is to explicitly model the molecular and mixture complexities and competitive interactions that underlie such data. Odotopes *per se* are not discussed in the present paper.

### The diversity of pharmacological interactions

Partial, weak, and antagonistic ligand-receptor interactions are fundamental features of basic pharmacology, and odorant receptors enjoy no exception from these physical principles. Indeed, interactions between related ligands and identified odorant receptors have been shown to exhibit diverse efficacies [32, 33, 43, 44]. As the chemical environment encountered by odorant receptors is largely unregulated, it has long been recognized that these nonlinear competitive interactions exert a substantial influence on peripheral odor encoding and contribute to the complexity of odor mixture interactions [34–36, 44, 47, 49, 84]. Contemporary studies revisiting or rediscovering this phenomenon have made broad measurements of odor and mixture effects upon OSN activation levels in the intact epithelium that reflect these pharmacological principles, illustrating that odorant receptor antagonism is a common and widespread phenomenon [37–39, 41, 42]. In particular, it has been observed that this antagonism has benefits for representing mixtures containing larger numbers of different odorants without saturation [39], as illustrated here in Figure 4. The improved “encoding capacity” that these authors suggest arises from antagonism, however, would not arise from sparseness – which is not directly relevant here – but for the reasons outlined in *Effects of different receptor efficacy Gaussian functions* and illustrated in Figure 4. Adherence to basic pharmacological laws also does not constitute an “evolutionary solution to the complexity of natural odors”. Similarly, the related claim that antagonism provides a form of normalization – specifically, rendering the distribution of responses in the OSN population independent of the number of components in a mixture [40, 42] – is in error. The error largely arises from the ad hoc model developed in [40], which discretizes various interactions and responses rather than employing established pharmacological laws. Figure 4 illustrates that it is receptor occupancy saturation, irrespective of the proportion of antagonistic responses, that produces the state in which additional ligands no longer systematically alter the mean activation level of the OSN population (i.e., where the activation curve becomes asymptotic; compare panels A-F, *middle panels*). Instead, the degree of antagonism (more specifically, the distribution of efficacies) influences the mean receptor activation level that is associated with this occupancy saturation (an effect roughly reflected by Figure 4A in [40]). The inclusion of antagonistic responses is nevertheless beneficial, though, as it improves individual OSNs’ capacity to respond to small changes in odor quality even after occupancy saturation.

As a general principle, the encoding of environmental odorant information by primary receptor activation patterns is heavily constrained by the physics of ligand-receptor interactions and should be understood as such. Biological adaptations to the problem of chemosensation are many and diverse – and may include details of OSN physiology such as nonzero baseline activation levels [85], which enable the phenomenon of inverse agonism [38, 85], and receptor reserve heterogeneity [56], which enables populations of sibling OSNs to encode broader ranges of ligand concentration. However, the unavoidable degeneracy of primary chemosensory sampling – in which levels of receptor activation cannot individually identify the ligands or combinatorial groups of ligands that evoke that activation – does not qualify as an adaptive “computation of mixture information”; rather, it should instead be considered as a problem requiring resolution by downstream neural circuitry [81].

Additional physiological and pharmacological effects exist that have been described or suggested for odorant receptors but are not explicitly modeled here: inverse agonists, inhibitory responses, and noncompetitive interactions. Inverse agonists are distinguished from (neutral) antagonists only when a receptor-coupled effector mechanism exhibits some nonzero level of constitutive activation. Under such circumstances, a ligand that, when bound, reduces effector activation below this baseline level qualifies as an inverse agonist. This effect can be measured as a simple negative efficacy [86], but in the present context this is a heuristic that obscures the nature of the interaction. To wit, the (negative) efficacy of such a ligand-receptor interaction is not a characteristic property of that interaction, as it also depends upon the level of and basis for the constitutive activity of the system against which it is measured. In the present framework, the efficacy parameter represents the proportion of potential receptor activation that a given ligand would evoke at full occupancy, and hence is limited to [0,1] on principle. If receptor-coupled effector activity has a constitutive level of, say, 20%, then a ligand exhibiting an efficacy of 0.2 at that receptor would be assessed as a neutral antagonist, whereas ligands exhibiting lower efficacies would be considered inverse agonists. Constitutive activity in OSNs is thought to depend on constitutive activity in ORs [85, 87], and as such could be modeled by adding one or more ligand points to reflect this background activity level and enable a functional result of inverse agonism. However, if effectors present a level of constitutive activity that is decoupled from receptor activation – potentially a much more complex circumstance – then this would be modeled in the present framework by adding relevant parameters describing this baseline activity and adjusting the activation function in the receptor objects to incorporate these new parameters.

Many inhibitory responses to odorants described to date in vertebrate OSNs appear to derive from inverse agonism [38, 85]. This must be distinguished from parallel OSN receptor-effector pathways that generate genuine inhibitory responses, as have been long established in lobster OSNs ([88]; see also [89]) and more recently demonstrated in mammalian OSNs [90–92], presenting the prospect of a form of opponent encoding [91]. In the present computational framework, the latter phenomenon would be most conveniently modeled by deploying multiple Gaussian pairs (receptors) that converge onto a given glomerulus object, one (or more) of which contribute negative activation to that object. An intermediate OSN object of course could be constructed if warranted. The efficacy parameter would remain in the [0,1] range for all ligand-receptor interactions.

Noncompetitive (including allosteric) receptor interactions [31, 36, 93] can be arbitrarily diverse, underlying inhibition, facilitation, and more complex interactions, and are not explicitly included in the present framework. If desired, these effects would be modeled using multiple Gaussian pairs converging on a given receptor object with the specific interactions of interest encoded into that receptor object.

### Summary

Odorant receptors obey established pharmacological laws as a matter of first principles. In comparison to other membrane receptors, they warrant special consideration only because of the relatively unregulated environment in which they are deployed. To wit, a cholinergic receptor gains that name because, within its regulated physiological environment, no significant agonist or antagonist other than acetylcholine is ever present; the receptor never naturally encounters the great diversity of other agonists, partial agonists, and antagonists that exist for it. In contrast, when odorant receptors are deployed into the olfactory epithelium, the diversity of ligand encounters is largely unregulated. The resulting degeneracy prevents specific interpretation of individual receptor activation. Specificity is gained instead by deploying large numbers of receptor types (hundreds in mammals) with diverse chemoreceptive fields into the olfactory epithelium. There, chemical ligands interact with multiple sensitive receptors, with individual interactions exhibiting characteristic affinities and efficacies. This combinatorial specificity provides information sufficient to counteract individual receptor degeneracy, and the peripheral olfactory system is structured to harness this information. However, the complexity of these many interacting pharmacological and neurophysiological variables renders unclear the quantitative properties of the resulting system and the outcome of any particular environmental sampling event. How does the peripheral olfactory system leverage the benefits and mitigate the limitations of the physical sampling principles imposed upon it?

Theoretical frameworks such as that presented here impose discipline on hypothesis generation when the systems under study are too complex to simply intuit [78, 94]. In the present framework, we describe a computational platform in which odorant ligands can be distributed in a physicochemical quality space and sampled by batteries of odorant receptors according to established pharmacological laws. This framework enables study of how mixture interactions and complex environments affect odor sampling, the impact of antagonistic and partial-agonist interactions on odor discrimination capabilities, and the utility of single-receptor convergence patterns onto olfactory bulb glomeruli. It also demonstrates the value of larger receptor deployments, and enables comparisons of how effectively different receptor complements (species-specific noses) sample a common physicochemical environment. In the present study, we focused on uniform distributions of receptors and ligands in order to establish the core properties of the system. Among other findings, we illustrate the likely basis for the phenomenon of “olfactory white” as a form of regression to the mean, and show that reducing the specificity of axonal convergence onto olfactory bulb glomeruli will systematically reduce the sensitivity of odor discrimination. Additional applications for this model framework include olfactory natural scenes studies based on nonuniform ligand distributions within Q-space and the adaptive properties of particular receptor deployments with respect to specific chemical environments, as well as assessment of the benefits and limitations of uniglomerular versus multiglomerular sampling by mitral cells.

## Acknowledgments

We gratefully acknowledge Michael Mariscal for advice on figure presentation. This work was funded by the National Institute on Deafness and Other Communication Disorders (R01 DC014701 to TAC and CRY).

